# Dissociable reactivation during NREM and REM sleep supports memory consolidation and emotional dissipation

**DOI:** 10.64898/2026.08.04.742295

**Authors:** Yuqi Zhang, Ziqing Yao, Danni Chen, Tao Xia, Lingqi Zhang, Andrew F. Luo, Xiaoqing Hu

## Abstract

Sleep is critical for memory consolidation and emotional regulation, yet the respective roles of non-rapid eye movement (NREM) and rapid eye movement (REM) sleep remain unclear. Here, using a within-subject crossover design, we recorded high-density electroencephalography (EEG) across two experimental nights while participants viewed neutral or aversive film clips in a counterbalanced order. Combining with daytime functional localizers establishing neural patterns of aversive vs. neutral emotional processing, multivariate pattern analysis revealed that the reactivation of aversive vs. neutral memory during nocturnal sleep was both stage-dependent and event-specific. In NREM sleep, valence-specific reactivation was time-locked to slow oscillation (SO)-spindle complexes but not to either event alone; in REM sleep, reactivation occurred selectively during phasic REM periods marked by rapid eye movements. Critically, NREM SO-spindle coupling percentage was associated with consolidation of temporal memories; whereas phasic REM reactivation strength was linked to overnight dissipation of negative affect. Our findings provide direct evidence that sleep reprocesses emotional experiences through dissociable stage- and event-specific mechanisms, laying out a framework for future targeted sleep-based interventions.

## Introduction

Memories of aversive or traumatic events are often more vivid and longer lasting than those of neutral experiences^1,2^. Although remembering aversive experiences can be adaptive by facilitating avoidance of future threats, it becomes maladaptive when emotional responses are overwhelming, resulting in pathological hyperarousal and involuntary intrusive memories^3–5^. Indeed, dysregulated emotional memory is characteristic of various psychiatric disorders, including post-traumatic stress disorder (PTSD), anxiety, and depression^6^. Sleep is recognized as a critical window for emotional memory re-processing^7,8^, with sleep-based interventions being highlighted as a promising approach for modifying maladaptive emotional memories bypassing conscious awareness^9–11^. However, the respective contributions of different sleep stages— particularly NREM and REM sleep—to emotional memory reprocessing, as well as their underlying neural mechanisms, remain poorly understood^7,12,13^. Clarifying these mechanisms not only contributes to fundamental understanding of memory consolidation and emotion regulation, but also bears substantial clinical implications in developing novel precision interventions for mood- and memory-disorders.

Theoretical accounts and emerging empirical evidence suggest that both NREM and REM sleep contribute to emotional memory re-processing yet in different roles^7,8^. NREM sleep, particularly slow-wave sleep (SWS), has been most consistently implicated in memory reactivation and systems-level consolidation of newly encoded memories^8,14,15^. Mechanistically, systems-level consolidation is mediated via coordinated coupling of cardinal neural oscillations during SWS, namely the 0.5-1.25 Hz cortical slow oscillations (SOs), 11-16 Hz thalamo-cortical sleep spindles, and 80-120 Hz hippocampal sharp-wave ripples (SWRs). During this orchestrated, cross-regional coupling, memory traces are reactivated and consolidated via hippocampal-neocortical communications, enabling long-term stabilization^16–18^.

Critically, SO-spindle coupling supports not only the consolidation of episodic details, but also the higher-order temporal structures of episodic events. In rodents, hippocampal neural activity patterns encoding prior waking experience are sequentially replayed during sleep, reinstating the temporal structure of recently encoded events that supports subsequent spatial navigation^19^. In humans, sleep has similarly been shown to enhance sequence memory, with SO-spindle coupling predicting the magnitude of these behavioral benefits^20,21^. Moreover, NREM sleep may adaptively transform emotional memories by differentially impacting content and affective tone of these memories^22–24^. Supporting this view, both mnemonic and affective responses to emotional experiences have been linked to NREM-related neural activities, including SWS duration, slow-wave EEG power, spindle activity, and, more recently, SO-spindle coupling^25–29^.

In parallel, REM sleep may specifically regulate the affective tones of emotional memories. Specifically, longer REM duration, greater REM continuity, and stronger REM theta activity have been linked to reduced next-day emotional intensity and attenuated amygdala reactivity to emotional experiences, whereas fragmented REM sleep appears to disrupt this adaptive emotional dissipation^30–36^. Yet REM sleep is far from homogeneous at finer temporal scales. Phasic REM periods—brief epochs characterized by bursts of rapid eye movements—are accompanied by transient cortical, limbic, and autonomic activation, setting them apart from the comparatively quiescent tonic REM periods^37,38^. Importantly, rapid eye movements themselves have been linked to amygdala activity and overt emotional behaviours during dreaming, suggesting that phasic REM may demarcate brief windows of emotion-related neural processing^39–42^. These observations raise the untested possibility that phasic REM provides a privileged window for emotional memory reinstatement, thereby facilitating adaptive emotional dissipation.

Indeed, to date, it remains unclear how NREM and REM sleep may directly engage in the reactivation of emotional memory, how they may differentially impact next-day memory and emotional responses, and whether stage-specific sleep neural activities may contribute to reactivation and behavioural impacts. Nevertheless, converging lines of evidence have begun to suggest that NREM and REM sleep may support emotional memory processing in complementary ways. For example, PSG-fMRI evidence indicates that SWS and REM sleep are associated with distinct neural signatures of negative memory consolidation, with SWS predicting better retention and reduced hippocampal responses, and REM sleep predicting stronger hippocampal-neocortical connectivity^43^. Shedding light on the potential causal contributions, stage-specific targeted memory reactivation studies showed that cueing emotional memories during NREM aids memory consolidation via spindle-related neural activity, whereas REM cueing may instead promote forgetting or affective downregulation^13,24,26,30^. Relatedly, a recent study employed an ensemble classifier to decode emotional patterns during sleep, revealing that the reprocessing of negative (vs. neutral) memories occurred across both NREM and REM stages, with the second sleep cycle emerging as particularly important for memory reactivation^44^.

Despite these advances, several notable methodological limitations constrain the strength of evidence. First, relatively few studies have directly compared stage-specific contributions within a single experimental framework^13,24^. Second, stage-specific accounts have often been inferred indirectly from changes in sleep macro-architecture or from a priori selected electrophysiological correlates, rather than from direct evidence of emotional memory reactivation during sleep^45^.

Third, most, if not all, studies use static images as stimuli, which inadequately capture the complexity and temporal structure of real-world emotional experiences^46^. These limitations obscure the precise mechanisms through which NREM and REM sleep contribute to emotional memory reprocessing.

Bridging these significant gaps, we investigated sleep-dependent emotional memory reactivation using an ecologically valid trauma film paradigm that simulates naturalistic aversive experiences^47–49^. In a within-subject crossover design, participants viewed either neutral or trauma film clips before sleep across two counterbalanced nights while high-density electroencephalography (EEG) was recorded throughout the nocturnal sleep. Critically, because the film clips unfolded as temporally structured naturalistic events, we assessed not only affective responses but also how sleep reshaped the temporal memories of emotional experience, moving beyond conventional item-based memory tests. We then combined multivariate decoding to test whether neural patterns distinguishing aversive from neutral experiences re-emerged during NREM and REM sleep stages and more specifically cardinal sleep oscillations and events. This decoding approach enabled us to move beyond indirect stage-level associations and directly characterize endogenous emotional memory reactivation at the level of discrete neural events^16^. Here, we found that emotional memory reactivation was detectable during both SO-spindle complexes and phasic REM periods, with these event-specific reactivation patterns differentially predicting overnight temporal memory consolidation and emotional dissipation.

## Results

### Overnight emotional dissipation and temporal memory consolidation

To characterize how sleep impacted affective and mnemonic outcomes, we examined state-level affect across the experimental session (aversive vs. neutral) and memory-specific emotional ratings during deliberate memory recall before and after sleep. For state-level affect, a 2 (session: aversive vs. neutral) × 4 (time: T0-T3) repeated-measures ANOVA revealed significant session × time interactions for state anxiety, *F*(2.16, 90.75) = 19.96, *p* < .001, 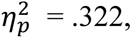 negative affect, *F*(1.98, 83.03) = 11.19, *p* < .001, 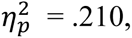 and positive affect, *F*(2.44, 102.66) = 14.62, *p* < .001, 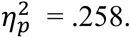 Post-hoc contrasts showed that the two sessions did not differ at baseline (T0). Following trauma exposure, participants in the aversive session exhibited elevated state anxiety and negative affect, together with reduced positive affect, both immediately after encoding (T1) and prior to sleep (T2). After sleep (T3), between-session differences in negative and positive affect were no longer detectable, whereas state anxiety remained elevated in the aversive session, indicating overnight attenuation of trauma-induced affective changes (Fig. 2A, full follow-up contrast statistics are reported in Supplementary Table 1). Memory-specific emotional ratings collected during pre- and post-sleep recall showed a similar pattern of emotional dissipation: significant session × time interactions emerged for recalled valence, *F*(1, 42) = 7.35, *p* = .010, 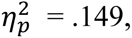 and traumatic impact, *F*(1, 42) = 8.16, *p* = .007,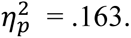 Pre-post contrasts confirmed that recalled memories were rated as less negative and less traumatic after sleep in the aversive session, but not in the neutral session. By contrast, arousal ratings showed no session × time interaction, *F*(1, 42) = 0.06, *p* = .801, 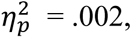 and declined similarly across both sessions overnight (full contrast statistics reported in Supplementary Table 2).

**Fig. 1.**
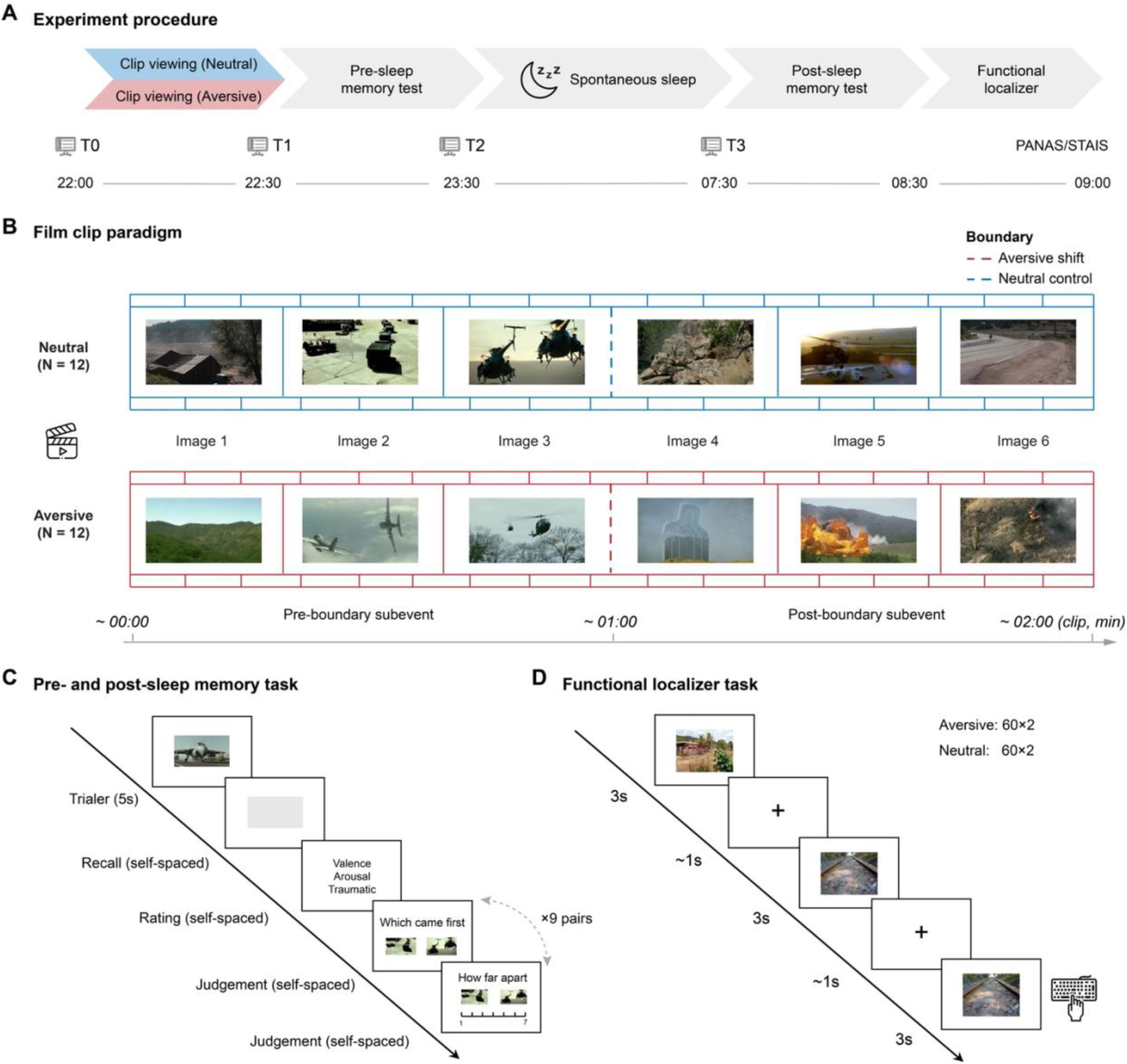
Experimental design. (**A**) Participants completed three laboratory visits: an adaptation night followed by two counterbalanced experimental nights, separated by one week. Each experimental session (aversive vs. neutral) comprised viewing 12 film clips, pre- and post-sleep memory assessments, overnight sleep recording, and an independent functional localizer task. (**B**) In each session, participants viewed either neutral or aversive film clips. Neutral clips contained exclusively neutral content throughout, whereas aversive clips began with a neutral lead-in followed by an aversive experience (all film images depict scenes without identifiable individuals and are shown for illustrative purposes only). (**C**) During the memory test, participants mentally recalled each event and rated their emotional responses to it. Temporal memory was assessed using image-pair comparisons drawn from three boundary-defined conditions: pre-boundary, post-boundary, and cross-boundary. Temporal-order judgments and temporal-distance estimations indexed memory for event structure. (D) In an independent functional localizer task, participants viewed aversive and neutral images selected from standardized affective image databases (see Methods for stimulus details) while performing a one-back working memory task.

**Fig. 2.**
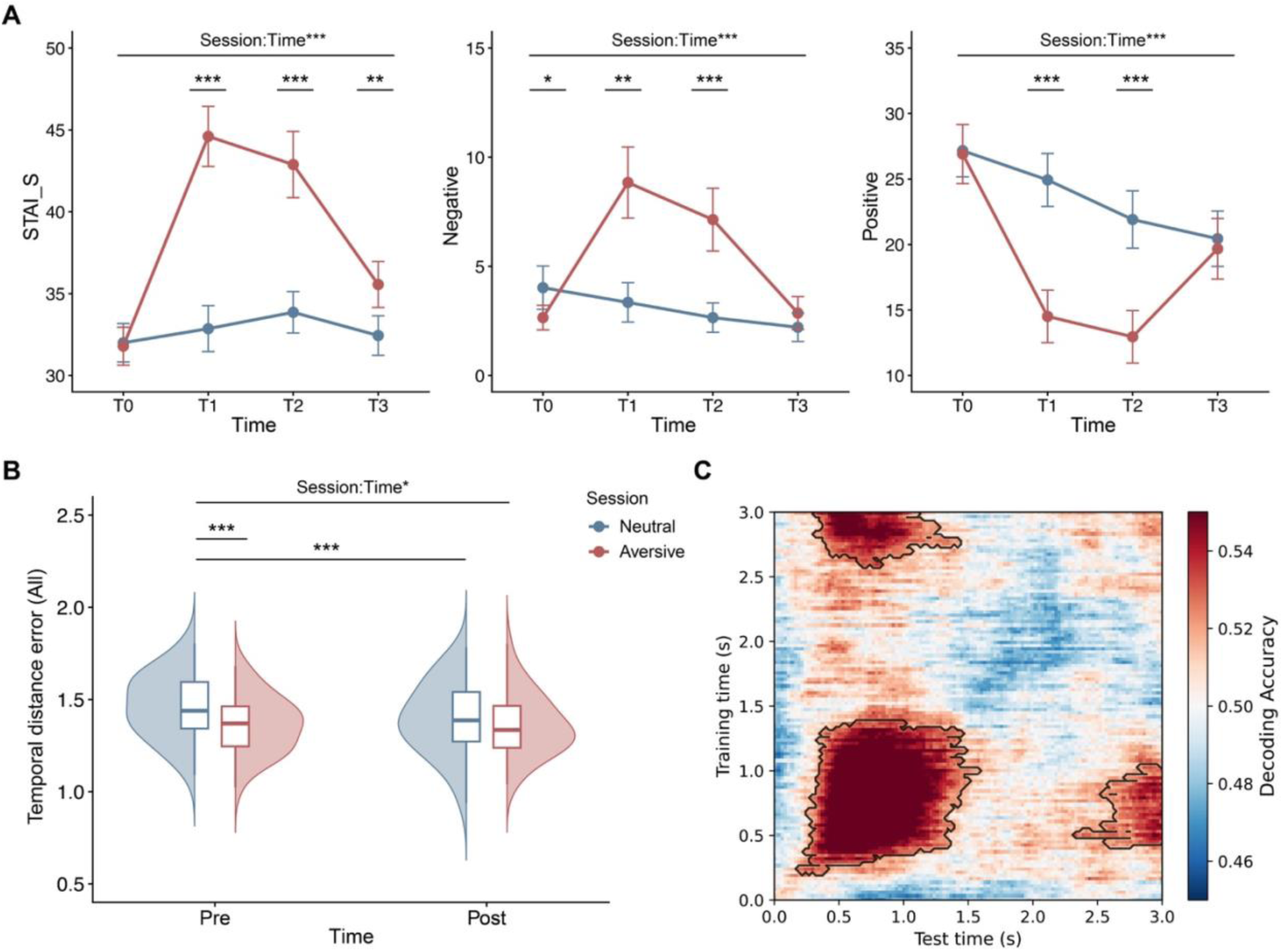
Behavioral results and wakeful valence-specific EEG decoding. (**A**) State anxiety, negative affect, and positive affect at baseline (T0), post-encoding (T1), pre-sleep (T2), and post-sleep (T3) in the neutral and aversive sessions. Trauma exposure increased state anxiety and negative affect and reduced positive affect after encoding; these effects were partially attenuated following sleep. (**B**) Temporal-distance estimation error before and after sleep. Temporal-distance error decreased after sleep in the neutral session, eliminating the pre-sleep session difference. (**C**) Wakeful valence-specific neural patterns. Temporal generalization decoding distinguished aversive from neutral images. Black contours indicate above-chance decoding (*p* < .05, cluster-corrected).

Regarding temporal memories, sleep did not affect temporal order accuracy (all *ps* ≥ .143). In contrast, sleep selectively improved temporal-distance memory for neutral, but not aversive, events (Fig. 2B), as indicated by a significant session × time interaction, *F*(1, 42) = 5.95, *p* = .019,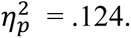 Before sleep, participants showed greater pre-sleep temporal precision for aversive than for neutral events, *t*(42) = 3.59, *p* < .001. After sleep, this session difference was no longer significant, *t*(42) = 1.27, *p* = .212. Pre–post contrasts further showed that temporal-distance error decreased significantly after sleep for neutral events, *t*(42) = 3.57, *p* < .001, but remained stable for aversive events, *t*(42) = 0.11, *p* = .912. Because emotional shifts in film clips can serve as salient event boundaries that may influence temporal memory^50–52^, we further examined whether sleep differentially affected temporal-distance memory for subevent pairs falling before, after, or spanning the onset of aversive content (i.e., the emotional boundary). Post-boundary pairs, which showed the largest emotional divergence between sessions, recapitulated the overall session × time pattern (Supplementary Note 1).

### Emotional valence is reliably decoded from the wakeful functional localizer

Prior work has shown that cardinal sleep events (e.g., slow oscillations, spindles) can carry memory-specific information (e.g., houses vs. faces) and support memory consolidation, but whether these events also reinstate the affective content of prior experiences remains unclear _16,18,53_. We therefore asked whether valence-specific representations could be identified in wakeful neural activity, and whether we could subsequently track valence-specific neural reactivation during sleep. To establish valence-specific neural patterns, we applied multivariate pattern analysis (MVPA) to EEG data recorded during a functional localizer task performed while awake, in which participants viewed aversive and neutral images and performed a one-back task (with 20% of images repeated) to maintain attention. A linear support vector machine (SVM) classifier reliably discriminated aversive from neutral trials in the wakeful localizer, with above-chance classification along the temporal generalization matrix diagonal from approximately 220 to 1500 ms after stimulus onset (Fig. 2C; see Supplementary Fig. 1 for the corresponding accuracy time course). Cross-time generalization further revealed a sustained, block-like pattern spanning roughly 250 to 1500 ms, with an additional significant window from 2500 to 3000 ms, indicating temporally extended valence-specific neural patterns (*p*_corrected_ < .05, Fig. 2C). Together, these analyses established a robust wakeful valence-specific neural template that was then used to track emotional memory reactivation during sleep.

### Emotional memory reactivation is specific to SO-spindle complexes and Phasic REMs

Applying this valence-specific classifier to sleep EEG data, we specifically examined slow oscillations (SOs), sleep spindles, and SO-spindle complexes during NREM sleep (Fig. 3A-B) and phasic REM periods (Fig. 4A-B) during REM sleep. Above-chance classification would indicate that neural activity surrounding a given sleep event contained valence-specific neural representations, and would be interpreted as reactivation of emotional memory during this particular sleep event.

**Fig. 3.**
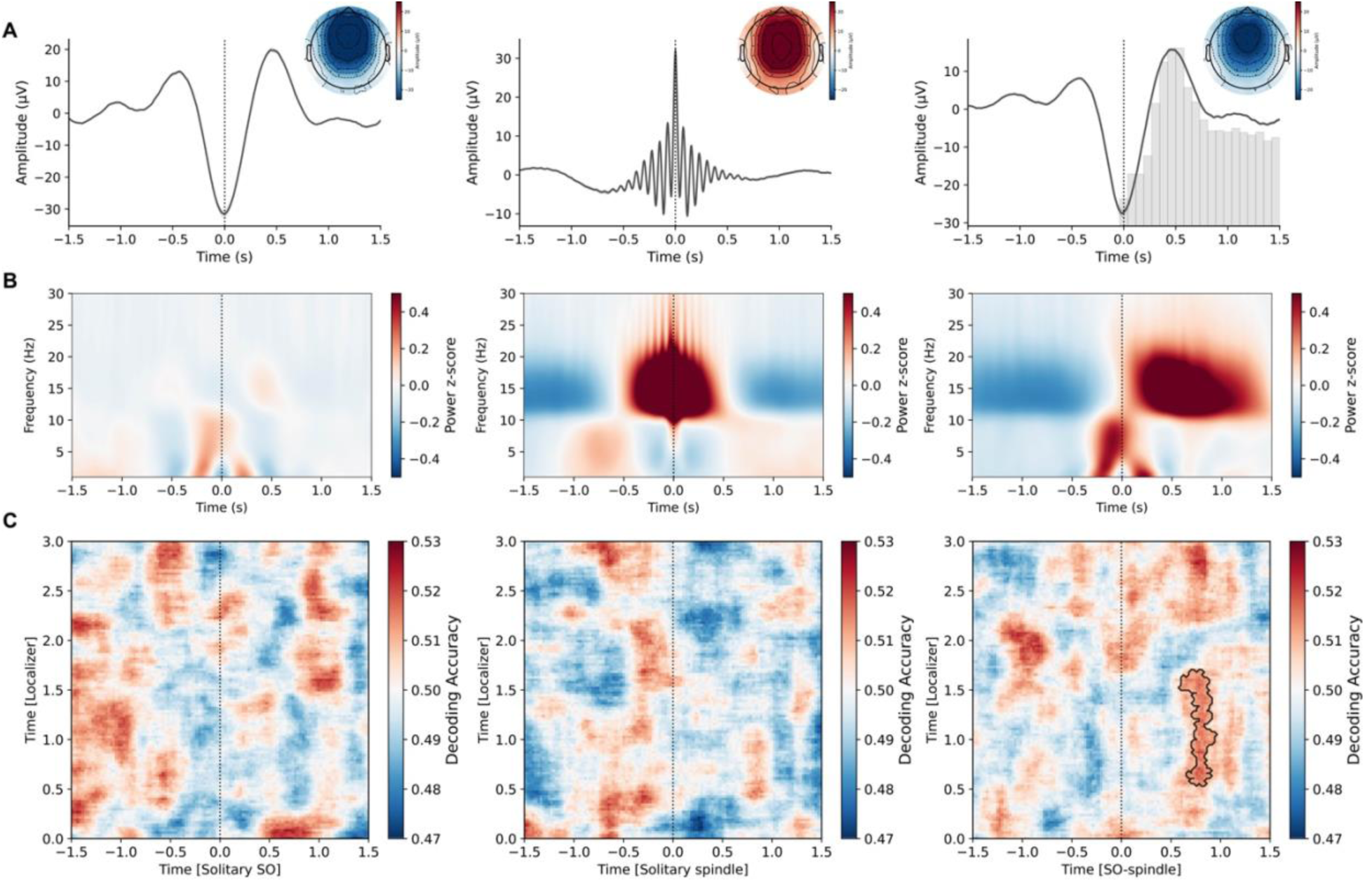
Emotional valence decoding during NREM sleep events. (**A**) Event-locked EEG waveforms for isolated slow oscillations (SOs; left), isolated spindles (middle), and SO–spindle complexes (right). Insets (top right) show the scalp topography of amplitude at the event center (t = 0). (**B**) Corresponding time-frequency representations for each NREM event type. (**C**) Event-locked valence classification. Classification accuracy did not exceed chance during isolated SOs (left) or isolated spindles (middle), but was significantly above chance during SO-spindle complexes (right), indicating that emotional valence information was selectively detectable during temporally coordinated SO-spindle events. Black contours indicate above-chance decoding (*p* < .05, cluster-corrected).

**Fig. 4.**
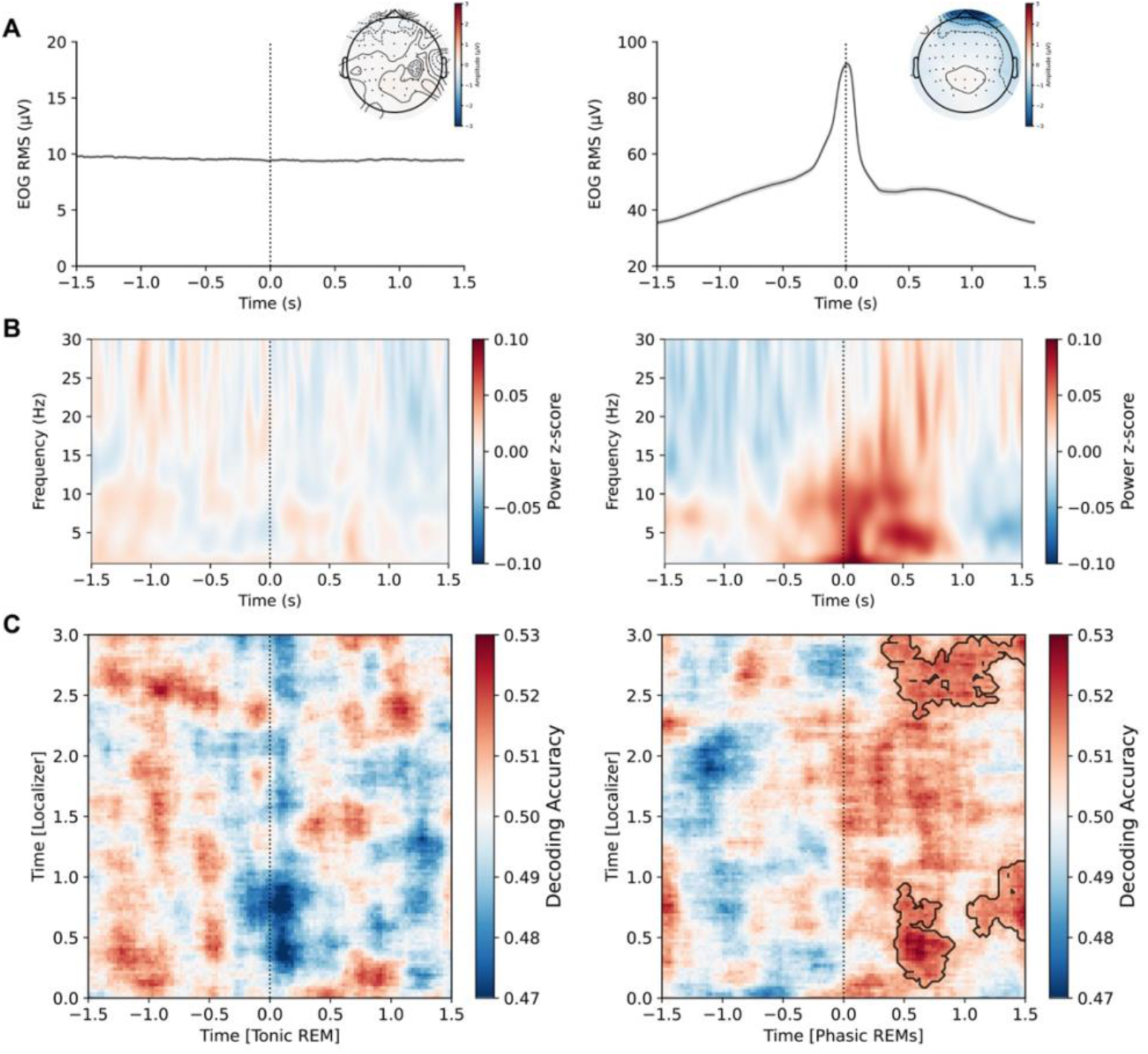
Emotional valence decoding during REM sleep events. (**A**) Event-locked EOG activity for tonic REM control periods (left) and phasic REM events (right). Insets (top right) show the scalp topography of amplitude at the event center (t = 0). (**B**) Corresponding time-frequency representations for each REM event type. (**C**) Event-locked valence classification. Classification accuracy did not exceed chance during tonic REM control periods (left), but was significantly above chance during phasic REM events (right), indicating that emotional valence information was selectively detectable during phasic REM. Black contours indicate above-chance decoding (*p* < .05, cluster-corrected).

Our results revealed emotional memory reactivation during both NREM and REM sleep. During NREM sleep, SO-spindle complexes showed significant above-chance decoding approximately 700-850 ms after the SO trough, around the emergence of spindles, with effects most evident for the 500-1600 ms localizer training time window (*p*_corrected_ <.05, Fig. 3C, right). In contrast, no robust above-chance decoding was observed during neither isolated SOs (Fig. 3C, left) nor isolated spindles (Fig. 3C, middle), indicating that NREM emotional memory reactivation was most evident during temporally coordinated SO-spindle events rather than during either isolated event. We then investigated whether individual differences in conventional SO-spindle coupling metrics would explain SO-spindle memory reactivation strength, yet we did not find significant results (preferred coupling phase, circular-linear *r* = .120, *p* = .787; coupling strength, Pearson’s *r* = .039, *p* = .830, 95% CI [−.308, .377]). Thus, although emotional memory reactivation was most evident among SO-spindle coupling events, participant-level coupling phase/strength did not account for individual differences in reactivation strength in the current study.

During REM sleep, reactivation was strongly associated with phasic REM periods characterized by burst of rapid eye movements. Significant above-chance decoding emerged 400-1500 ms after the REM peak, primarily for localizer training windows spanning 100-1000 ms and 2500-3000 ms (*p*_corrected_ <.05, Fig. 4C, right). To test whether this effect was specifically tied to phasic REM rather than REM sleep more broadly, we randomly sampled matched tonic REM periods in which rapid eye movements were absent and used them as control events (Fig. 4A-B, left). Phasic REM events showed widespread increases in delta, theta, alpha, and beta power relative to tonic REM events (Supplementary Fig. 2), consistent with a neurophysiologically distinct phasic REM microstate. Critically, no significant decoding was observed during tonic REM control periods (Fig. 4C, left), indicating that emotional memory reactivation was specific to phasic REM microstates. Given that phasic REM is characterized by eye movements, we next assessed the contribution of eye-movement-related signals to phasic REM decoding. We applied independent component analysis (ICA) to identify eye movement-related components that were dominated by ocular activity. Significant phasic REM decoding persisted after removal of these ocular-dominated components. Conversely, decoding based only on the ocular-dominated components did not yield above-chance classification (Supplementary Fig. 3). These patterns strongly indicate that the phasic REM decoding effect was not driven by ocular signals alone, but was primarily driven by cortical EEG activity associated with phasic REM. Together, these findings indicate that emotional memory reactivation during sleep was not uniformly distributed across sleep stages, but emerged preferentially during discrete neural events: SO-spindle complexes during NREM sleep and phasic REM microstates during REM sleep.

To further delineate the temporal dynamics of sleep-dependent emotional memory reactivation, we conducted a segment-wise decoding analysis for NREM SO-spindle complexes and phasic REM events. Sleep events were divided into five consecutive 90-min temporal segments, approximating the typical duration of a sleep cycle^53^, to characterize whether the strongest reactivation effects were concentrated in specific portions of the night. The first segment began 5 min after stable sleep onset to avoid the transitional period of unstable sleep (segment durations are reported in Supplementary Tables 3). For each event type and segment, we applied the same localizer-to-sleep decoding pipeline used in the main analysis. Because the number of sleep events varied across participants, sessions, and segments, a participant was included in a given analysis only if both aversive and neutral sessions contained at least 30 sleep events for that specific event type and segment. Group-level decoding was performed only when at least 12 valid participants were available (trial statistics are provided in Supplementary Table 4). This analysis revealed significant reactivation during the second segment for SO-spindle events and during the fourth segment for phasic REM events, suggesting temporally dissociable windows for mnemonic and affective reprocessing (Supplementary Fig. 4).

### SO-spindle coupling percentage predicts overnight temporal memory consolidation

To examine the functional significance of emotional memory reactivation during sleep, we first assessed whether reactivation strength predicted overnight changes in temporal memory performance. For each participant, reactivation strength was quantified as the mean decoding accuracy extracted from statistically significant clusters during SO-spindle complexes and phasic REM events. However, reactivation strength per se was not significantly associated with temporal memory changes for either event type (Supplementary Table 5), possibly because the emotion-specific classifier was optimized to detect emotional rather than memory content.

We next asked whether the prevalence of SO-spindle complexes, indexed by SO-spindle coupling percentage, would predict temporal memory consolidation. Indeed, using a linear mixed model with post-sleep temporal-distance error as the dependent variable and pre-sleep error as a covariate, we found a significant main effect of coupling percentage, *F*(1, 40.28) = 4.66, *p* = .037, indicating that higher coupling percentage was associated with lower post-sleep error (Fig. 5A). The coupling percentage × session interaction was not significant, *F*(1, 36.92) = 0.99, *p* = .327, suggesting that this association did not reliably differ between aversive and neutral sessions. Given that the behavioral benefits were most evident for neutral events, exploratory simple-slope estimates were examined for descriptive purposes. These showed a significant negative association in the neutral session, *z* = −2.20, *p* = .028, and a weaker, non-significant association in the same direction in the aversive session, *z* = −1.01, *p* = .313.

**Fig. 5.**
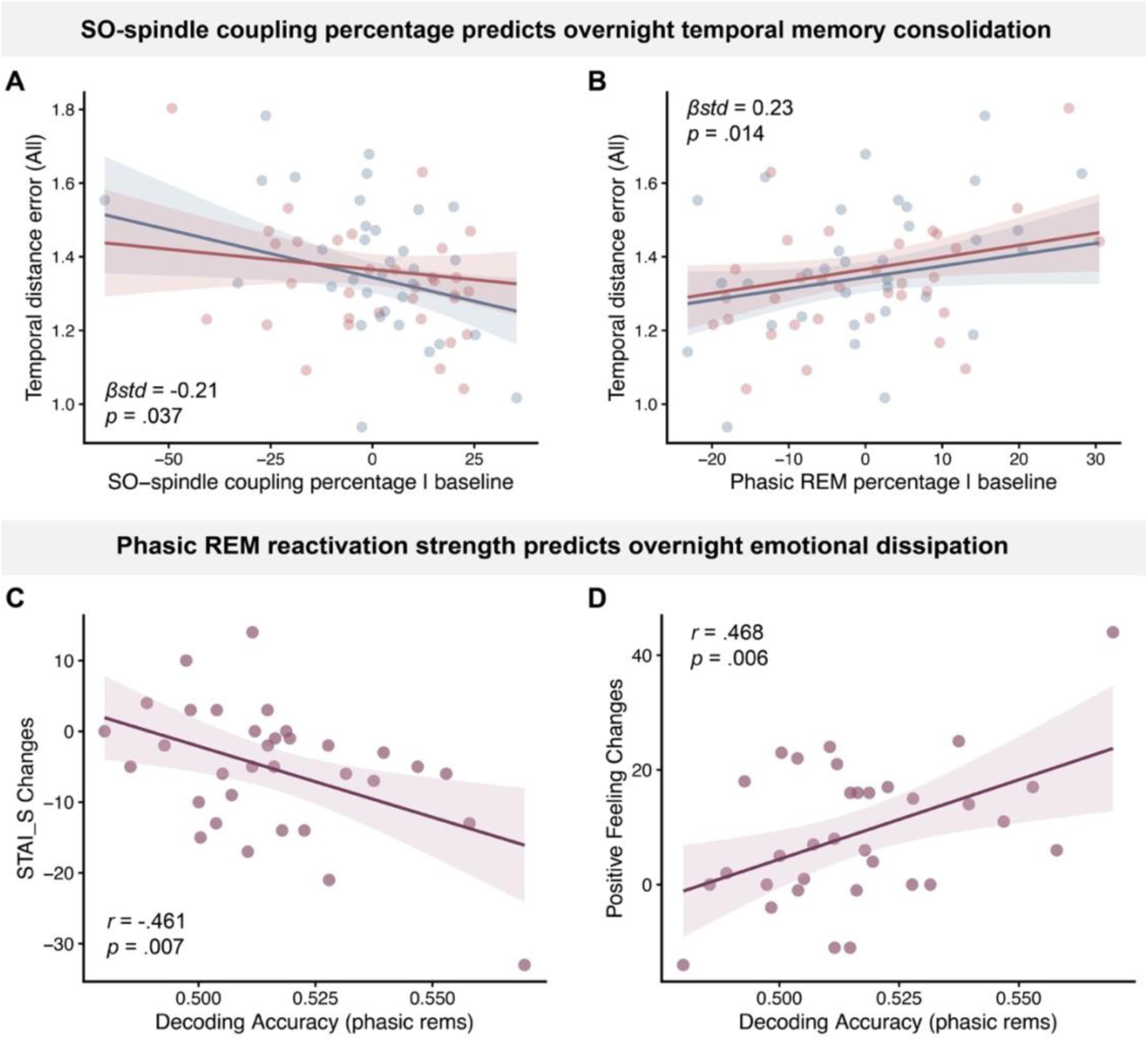
SO-spindle coupling percentage supports temporal memory consolidation, whereas phasic REM reactivation supports emotional dissipation. (**A**) Greater SO-spindle coupling percentage predicted lower post-sleep temporal-distance error (blue, neutral session; red, aversive session). (**B**) In contrast, greater phasic REM percentage predicted higher post-sleep temporal-distance error. Greater phasic REM reactivation strength was associated with larger overnight reductions in state anxiety (**C**) and (**D**) larger increases in positive affect.

**Fig. 6.**
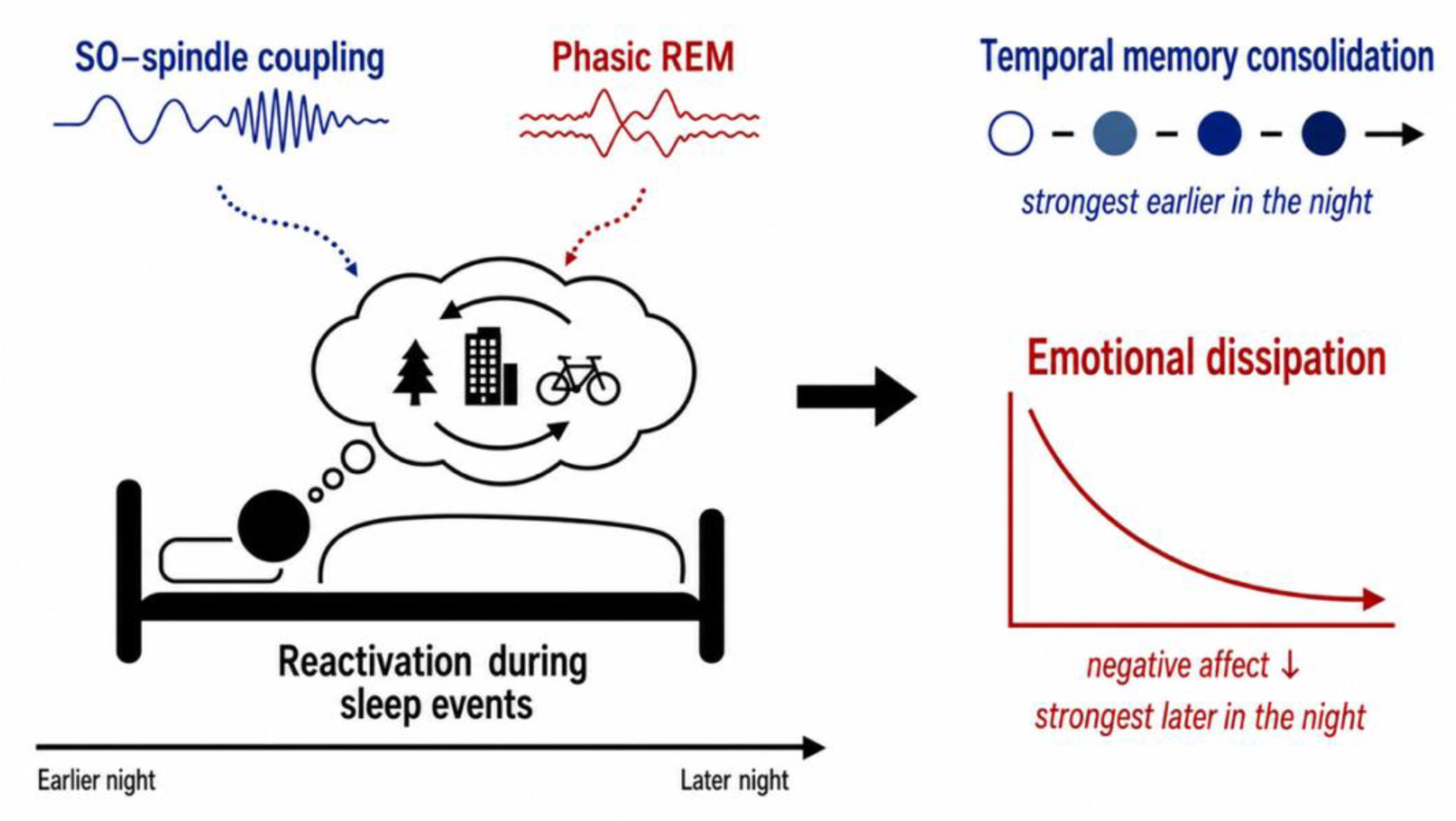
Dissociable roles of NREM and REM sleep in emotional experience reprocessing. Daytime experiences may be reactivated during cardinal sleep events for subsequent reprocessing. SO-spindle coupling, strongest earlier in the night, was linked to temporal memory consolidation, whereas phasic REM, strongest later in the night, was linked to emotional dissipation.

In contrast, phasic REM percentage provided no evidence for a memory benefit. Rather, higher phasic REM percentage was associated with greater post-sleep temporal-distance error after controlling for pre-sleep performance, *F*(1, 45.99) = 6.49, *p* = .014, with no interaction with session, *F*(1, 38.31) = 0.01, *p* = .932 (Fig. 5B). Phasic REM density showed the same pattern, with higher density predicting larger post-sleep error, *F*(1, 44.91) = 4.30, *p* = .044, and again no session interaction, *F*(1, 37.87) = 0.02, *p* = .887. Together, these findings indicate that the overnight temporal-distance memory benefit was specifically linked to coordinated SO-spindle events rather than to the overall prevalence of phasic REM sleep. Boundary-specific analyses are provided in Supplementary Note 3.

### Phasic REM reactivation strength predicts overnight emotional dissipation

We next examined whether event-specific reactivation strength predicted overnight changes in state and memory-specific affective responses. For memory-specific ratings, reactivation strength during neither SO-spindle complexes nor phasic REM events was significantly associated with overnight changes, all *ps* > .148 (Supplementary Table 6). For state-level affect, SO-spindle reactivation strength was unrelated to overnight changes in state anxiety, negative affect, or positive affect, all *ps* > .335. In contrast, phasic REM reactivation was robustly associated with overnight emotional dissipation. Higher reactivation strength time-locked to phasic REMs predicted larger overnight reductions in state anxiety, *r* = -.46, *p* = .007, *p_FDR_* = .010, 95% CI [-.69, -.14], and negative affect, *r* = -.44, *p* = .011, *p_FDR_* = .011, 95% CI [-.68,-.11], as well as greater increases in positive affect, *r* = .47, *p* = .006, *p*_FDR_ = .010, 95% CI [.15, .70], Fig. 5C-D.

We next examined the specificity and robustness of this relationship. First, phasic REM reactivation strength was not significantly correlated with pre-sleep affective differences between the aversive and neutral sessions in state anxiety, *ρ* = .260, *p* = .144, negative affect, *ρ* = .250, *p* = .160, or positive affect, *ρ* = -.236, *p* = .186; all *p_FDR_* > .185. Thus, stronger reactivation in phasic REM did not simply reflect greater pre-sleep emotional responses that carried over to sleep. Second, because rapid eye movements are a defining feature of phasic REM, we repeated the analyses after ICA-based removal of components dominated by ocular activity. The associations between phasic REM reactivation and overnight reductions in state anxiety, *r* = -.49, *p* = .004, *p_FDR_* = .013, 95% CI [-.71, -.17], and negative affect, *r* = -.42, *p* = .014, *p_FDR_* = .021, 95% CI [-.67, -.10], remained significant. The association with increased positive affect was attenuated and no longer significant, *r* = .31, *p* = .075, *p_FDR_* = .075, 95% CI [-.03, .59] (Supplementary Fig. 5). These results suggest that the links between phasic REM reactivation and reductions in negative affect were driven by cortical EEG activity associated with phasic REM. To further characterize the neurophysiological profile of phasic REM reactivation, we tested whether the decoding effect depended on any single canonical frequency band. After selectively removing delta (0.5-4 Hz), theta (4-8 Hz), alpha (8-12 Hz), or beta (13-30 Hz) activity from both the wakeful localizer and sleep-event data, phasic REM decoding remained above chance and its association with overnight emotional dissipation was preserved across all bandstop analyses (Supplementary Fig. 6). Thus, phasic REM reactivation was not attributable to any single frequency band, but instead reflected a broader EEG pattern associated with phasic REM microstates. Third, because trauma-analogue exposure altered sleep macrostructure (Supplementary Note 2), we conducted partial correlation analyses controlling for total sleep time, sleep-onset latency, REM proportion, and sleep efficiency. The associations between phasic REM reactivation and overnight emotional changes remained significant both when reactivation strength was estimated from the original EEG data and when it was re-estimated after ICA-based removal of ocular-dominated components (Supplementary Table 7), indicating that the phasic REM-driven emotion dissipation was not attributable to global sleep architecture differences. Finally, phasic REM reactivation was not significantly associated with depressive symptoms, trait anxiety, thought control strategies, empathy, or emotion regulation strategies, all *p_FDR_* > .05 (Supplementary Table 8), suggesting that this effect was not explained by stable individual differences in mood or emotion-related traits. Together, these findings support a specific link between phasic REM, emotional memory reactivation, and overnight emotional dissipation, particularly reductions in negative affective emotional states.

## Discussion

How does a night of sound sleep help resolve a distressing experience, while preserving the content of the memory? Using an independent wakeful valence localizer, we show that emotional memory reactivation during sleep is organized around specific microevents rather than uniformly expressed across the night. Valence-specific neural patterns preferentially re-emerged during SO-spindle complexes in NREM sleep and during phasic REM sleep, revealing two temporally precise, event-specific windows for emotional memory reactivation. Critically, these microevent-specific reactivations bear dissociable functional significance: the proportion of SO-spindle complex was associated with overnight changes in temporal memory, whereas phasic REM reactivation predicted overnight dissipation of negative affect. These findings reveal complementary and dissociable NREM and REM event-specific mechanisms through which sleep reshapes the mnemonic structure and affective tone of emotional experiences.

Although REM sleep has been regarded as instrumental for adaptive regulation of daytime stressful experiences, our NREM findings revealed that emotional memory reactivation also emerges within precisely coordinated SO-spindle complexes. NREM sleep oscillations (SOs, spindles, and ripples) have long been implicated in systems-level memory reactivation and consolidation, with particular emphasis on their temporal synchronization^8,18^. Our results extend prior work linking SO-spindle coupling to endogenous memory reactivation by showing that SO-spindle coupling can carry valence-specific neural patterns, not only declarative memory content^16,17,44,53^. It shall be noted that the strength of this SO-spindle memory reactivation did not predict overnight changes in either temporal memory or affective responses. This could be due to the specificity of our wakeful functional localizer that was trained solely to distinguish negative from neutral valence. Thus, the reactivation strength may primarily indicate reactivation of emotional content rather than the temporal structure of the aversive event memories. Thus, while SO-spindle complexes may support the reinstatement of emotional content, overnight changes in temporal memory may depend more on the degree of SO-spindle coupling than on emotion-specific reactivation strength itself. Indeed, participants with a higher proportion of SO-spindles coupling showed reduced temporal distance errors overnight, i.e., consolidation of temporal memories. This finding is consistent with prior work suggesting that SO-spindle coupling supports the consolidation of structured and sequential information—including linguistic sequences and real-world episodic memories—beyond retention of individual memory items^20,21^. Notably, the association between SO-spindle coupling and temporal memory consolidation was most evident for neutral events, mirroring the behavioral pattern in which neutral events showed the clearest overnight improvement in temporal memory. By contrast, aversive events already showed better temporal-distance memory before sleep, suggesting that their temporal organization may have been more strongly encoded or less malleable, leaving less room for additional sleep-dependent consolidation. This interpretation aligns with evidence that emotional arousal enhances memory encoding and consolidation^54,55^, can bias the organization of event memories^50,56,57^, and may leave weaker or more labile memory traces more susceptible to sleep-dependent benefits^58–61^.

Whereas NREM sleep was closely tied to SO-spindle coupling and temporal memory consolidation, REM sleep revealed a distinct pattern: emotional memory reactivation emerged selectively during phasic REM and predicted emotional dissipation rather than memory consolidation. This finding aligns with longstanding theories implicating REM sleep in affective regulation ^7,34^. Neurobiologically, REM sleep provides a unique context for emotional memory reprocessing, characterized by markedly reduced noradrenergic tone alongside sustained engagement of amygdala-hippocampal networks that encode salient memories^7,23^. Within this framework, REM sleep is thought to attenuate emotional intensity while preserving the memory trace itself. Consistent with this view, human studies have provided correlational evidence linking the amount of REM sleep to reduced next-day amygdala reactivity and subjective emotional intensity, whereas REM fragmentation appears to disrupt this adaptive process^33,35^. Moreover, causal evidence comes from REM targeted memory reactivation research, which shows that cueing negative memories during REM reduces subsequent amygdala and salience-network responses, physiological arousal, and subjective distress^30^. Our findings demonstrated that emotional memory reactivation and adaptive dissipation may happen specifically within brief phasic REM microevents rather than uniformly distributed across REM sleep. Indeed, phasic and tonic REM are increasingly recognized as functionally distinct microstates. Phasic REM is characterized by transient cortical and autonomic surges alongside diminished responsiveness to external stimuli, whereas tonic REM sustains a greater capacity for environmental monitoring^37,38,62^. Critically, rapid eye movements are time-locked to activation in limbic and medial temporal structures implicated in emotional processing—most notably the amygdala and hippocampus—as evidenced by both fMRI and intracranial recordings^39,63,64^. Phasic REM has therefore been hypothesized to facilitate the processing of emotional experiences, with important implications for emotion regulation and fear extinction^41^. By demonstrating that phasic REM reactivation strength predicted overnight emotional dissipation, our results provide direct evidence that these transient REM microevents may serve as privileged temporal windows for sleep-dependent emotional memory reprocessing.

Our control analyses further clarify the nature of phasic REM reactivation. The association between phasic REM reactivation and emotional dissipation persisted after ICA-based removal of eye-movement-related components, indicating that this effect reflected neural processes beyond ocular activity per se. Importantly, this does not diminish the potential significance of rapid eye movements themselves. Rather, REMs may serve as physiological markers—and potentially as organizing signals—of transient phasic microstates during which emotional representations are preferentially reprocessed. Consistent with this interpretation, phasic REM periods showed widespread increases in spectral power relative to tonic REM^37^. Moreover, removing individual frequency bands did not abolish either the decoding effect or its behavioral association, suggesting that phasic REM reactivation reflected a broadband neural signature rather than activity in any single canonical frequency band^53^. Together, these findings suggest that rapid eye movements index phasic REM microstates characterized by heightened cortical activation and limbic-autonomic engagement, thereby providing a favorable neurophysiological context for emotional memory reprocessing.

Although our daytime valence localizer enabled decoding of emotion-specific memory reactivation and revealed its dissociable affect and memory consequences, important questions remain for future investigation. First, although our study behaviorally characterized the temporal structure of emotional experiences, our decoding approach was optimized to detect valence-specific reinstatement rather than temporally unfolding reactivation or replay. As such, it cannot directly establish whether emotional memories are replayed in their original temporal order during sleep. Addressing this question will require sequence-sensitive decoding or representational similarity analyses, which could reveal how emotional memories unfold across sleep and how this unfolding shapes both the content and temporal organization of subsequent experiences. Second, although ICA-control analyses suggest that phasic REM decoding was not driven by ocular activity, scalp EEG does not allow precise localization of the underlying neural sources. Future work combining sleep EEG with fMRI or intracranial recordings could directly pinpoint the contribution of limbic and medial temporal circuits to phasic REM reactivation and emotional dissipation. Finally, although the trauma-film paradigm provides a more naturalistic and ecologically valid approach, it remains an analogue model. It may not fully capture the complexity, severity, personal relevance, or clinical persistence of real-world traumatic memories. Examining these sleep-dependent reprocessing mechanisms in clinical or subclinical populations will therefore be critical for establishing their relevance to maladaptive emotional memory processing and trauma-related disorders.

Overall, our study argues against a unitary account of sleep-dependent emotional memory consolidation. Instead, it supports a model in which distinct sleep events differentially shape the affective and mnemonic consequences of prior experience. By identifying these event-specific signatures, the present findings provide a framework for understanding how sleep adaptively reshapes emotional memories and for developing targeted sleep-based interventions.

## Methods

### Participants

Sixty healthy participants were recruited for a three-night sleep study. Seventeen participants were excluded from final analyses: eight withdrew after the adaptation night, seven withdrew after the first experimental night, and two were excluded due to careless responding on the behavioral task. The final behavioral analyses included 43 participants (mean age = 24.37 ± 3.01 years; 34 females; Supplementary Table 9 for demographic characteristics). Among these 43 participants, 33 participants were included for the overnight sleep EEG analyses across both experimental nights. The remaining ten participants were included in the behavoural analyses but were excluded from EEG analyses because of technical issues, including loss of M1/M2 reference channels, excessive electrode failure, or incomplete data saving. The sample size of the current study was comparable to, or larger than, those reported in previous human sleep studies (N = 32-32)^16,53,65^. For pre-screening, participants completed the Pittsburgh Sleep Quality Index (PSQI)^66^, the Beck Depression Inventory-II (BDI-II)^67^, and answered questions assessing general health, sleep habits, medication status, and recent life events. All included participants reported good sleep quality, no night-shift work within two weeks prior to the study, no current medication use, and no history of neurological or psychiatric disorders. The study was approved by the Human Research Ethics Committee of the University of Hong Kong, and all participants provided written informed consent before participation.

## Materials

### Film clips

We selected 12 trauma-analogue and 12 theme-/content-matched neutral film clips from online sources and prior trauma-film studies^68,69^. Each clip lasted approximately 2 minutes—longer than in previous studies—to enhance immersion and elicit strong aversive emotions, while allowing participants to engage fully with the film narratives^48,69,70^. Neutral clips were carefully chosen to match the trauma-analogue clips in setting and semantic content but featured distinct plots without distressing or aversive elements. All clips underwent identical video filtering to minimize differences in visual quality and color between trauma and neutral clips. Six independent raters evaluated each clip for emotional feelings, semantic content, visual and auditory complexity, and storyline complexity (see Supplementary Table 10 for details). As intended, trauma clips elicited significantly stronger negative affect than neutral clips, reflected in lower valence (*M* = 2.76 vs. 5.01, *p* < .001), higher arousal (*M* = 6.00 vs. 2.36, *p* < .001), and greater traumatic impact (*M* = 6.39 vs. 1.07, *p* < .001). Crucially, the two clip types did not differ in semantic content ratings (both *M* = 3.68, *p* = 1.000), confirming successful matching along these non-emotional dimensions. Visual complexity was rated slightly higher for trauma clips (*M* = 2.86 vs. 2.44, *p* = .019), whereas storyline and auditory complexity did not differ (both *ps* > .108). These results confirm effective manipulation of emotional intensity with well-controlled non-emotional features.

### Temporal memory pictures

To assess temporal memory for each film clip, six still frames were extracted at evenly spaced time points spanning the full 2-min duration of each clip. These frames provided a representative sample of each clip’s temporal progression. For neutral clips, all six frames depicted neutral content. For trauma clips, which typically consisted of a neutral lead-in followed by an aversive subevent, the first three frames depicted neutral content, and the next three frames depicted aversive content. This transition from neutral to aversive content was treated as an event boundary, allowing temporal memory performance to be examined separately for pre-boundary, post-boundary, and cross-boundary frame pairs. The extraction timepoints were matched between trauma and neutral clips across all six temporal positions, with no significant timing differences between conditions (all *ps* > .20; Supplementary Table 11). This ensured that differences in temporal memory performance could not be attributed to systematic differences in the sampling timepoints.

### Functional Localizer pictures

To train a functional localizer based on aversive vs. neutral EEG patterns, we used 240 pictures (120 aversive, 120 neutral) that were selected from internet affective picture databases, such as the International Affective Picture System (IAPS) and the Nencki Affective Picture System (NAPS). These databases provide normative ratings for valence and arousal, as well as semantic category labels. To approximate the content of the trauma and neutral film clips, images were selected primarily from two broad semantic categories: people and scenes. All images were standardized to landscape orientation and resized to 640 × 480 pixels to minimize low-level visual differences across stimuli. As expected, the selected negative images had significantly lower normative valence ratings (*M* = 1.93 vs. 4.86, *p* < .001) and significantly higher arousal ratings (*M* = 5.44 vs. 3.18, *p* < .001) than neutral images (Supplementary Table 12).

## Procedures

### Overview

Participants spent three nights in the sleep laboratory over two weeks. The first night served as an adaptation night to help participants adapt to the environment of the sleep lab. The two experimental nights were scheduled one week apart to minimize carryover effects, with aversive vs. neutral session order being counterbalanced across participants. On each experimental night, participants arrived at approximately 8:30 PM and completed baseline assessments of vigilance and emotional state before the film viewing task. After film viewing, emotional state was reassessed, followed by a pre-sleep temporal memory assessment. Participants then had an 8-hour sleep opportunity while 64-channel EEG was recorded. The following morning, approximately 25 min after awakening, participants completed post-sleep assessments of vigilance, emotional state, and temporal memory. A wakeful functional localizer was then administered to identify neural patterns associated with emotional processing. After each experimental night, participants completed a five-day online intrusion diary to track intrusive memories related to the film clips.

### Questionnaires and emotional calibration task

During the adaptation night lab visit, participants completed standardized questionnaires assessing subjective sleep quality, chronotype, depressive symptoms, prior trauma exposure, and emotion regulation styles. These included the Pittsburgh Sleep Quality Index (PSQI)^66^, the reduced version of the Morningness-Eveningness Questionnaire (rMEQ)^71^, the Beck Depression Inventory-II (BDI)^72^, the State-Trait Anxiety Inventory (STAI)^73^, the Trauma Events Questionnaire (TEQ)^74^, the Thought Control Questionnaire (TCQ)^75^, the Chinese version of the Interpersonal Reactivity Index (IRI-C)^76^, and the Emotion Regulation Questionnaire (ERQ)^77^. Participants then completed an emotional calibration task to verify that they could differentiate aversive from neutral film stimuli. Two short film clips, one trauma and one neutral, were each presented twice in randomized order. After each viewing, participants rated their emotional responses (valence, arousal, perceived traumatic impact) using the same measures as in the main experiment. Participants who failed to show sufficient differentiation between the trauma and neutral film clips, defined as highly similar ratings across conditions, were excluded from further participation.

### Psychomotor vigilance task (PVT)

Participants completed a PVT before sleep and after waking up to assess their vigilance levels. In each trial, a fixation cross was shown on the screen, followed by a counter that started counting up from 0 to 2s. Participants were instructed to press the space bar as quickly as possible to stop the counter once it appeared. The inter-trial interval ITI varied randomly between 2 and 10s. After each response, participants received feedback on their reaction time. The task lasted for 5 minutes.

### Emotional state ratings

Participants reported their current emotional state using a 10-point visual analogue scale (VAS) at four time points throughout the experimental session: T0 (baseline), T1 (after film-viewing), T2 (pre-sleep), and T3 (post-sleep). Positive affect items were adapted from the Positive and Negative Affect Schedule (PANAS)^78^, and included: Interested, Inspired, Relaxed, Enthusiastic, Content, and Happy. Negative affect items were drawn from a previous study employing a similar trauma film paradigm^79^, and included: Sad, Hopeless, Fearful, Anxious, Depressed, and Distressed. Participants rated their feelings on the VAS to indicate how they felt at each timepoint. Furthermore, participants completed a 20-item State-Trait Anxiety Inventory (STAI-S) to assess their current state of anxiety. This scale includes items such as “I feel tense” and “I am worried”, as well as reverse-coded items like “I feel calm” and “I feel secure”. Each item is rated on a 4-point Likert scale, ranging from “Almost Never” to “Almost Always”, with higher scores indicating greater state anxiety.

### Film viewing task

During the aversive session, participants viewed a series of 12 film clips depicting traumatic events such as car accidents, plane crashes, and shootings. Each trial began with a fixation cross on a black screen for approximately 1.5s, followed by the presentation of the film clip. Participants were instructed to watch each clip attentively for a later memory test. After each clip, participants rated their emotional responses using a 9-point Likert scale, assessing valence, arousal, and perceived traumatic impact. They also rated their familiarity with the content of the clip. The neutral session followed the same procedure, except that participants viewed neutral film clips depicting everyday scenarios, with semantic content being carefully matched to the aversive clips but devoid of emotional or traumatic elements. The inter-trial interval (ITI) between clips jittered between 4 and 6s. To ensure the experimental conditions are controlled, all film clips are presented in pseudorandomized order with no more than two scenes of the same type of scenes (e.g., car crash, gunshot) appearing consecutively.

### Pre- and post-sleep memory test

After watching the film clips, participants completed pre-sleep emotional and memory tests. Regarding the temporal memory, participants completed temporal order judgment and temporal distance estimation. Specifically, in each trial, participants were presented with a 5s segment depicting the first 5 s of the film clip as a reminder. Following the reminder, participants were instructed to mentally replay the full content of the clip in chronological order within a 2-minute time limit. After mental replay, they rated their emotional responses to the recalled content (valence, arousal, and perceived traumatic impact). Subsequently, participants were presented with each of nine pairs of screenshots extracted from the clip. For each pair, participants first indicated which of the two screenshots had appeared earlier in the clip (temporal order judgment), and then estimated the perceived temporal distance between the two screenshots on a 7-point scale (1 = very close in time, 7 = very far apart in time). The nine screenshot pairs were systematically constructed from the six screenshot images extracted from each clip at fixed temporal intervals. These pairs were categorized into three conditions based on their position relative to the within-clip boundary: (1) pre-boundary pairs (images 1-2, 2-3, and 1-3), (2) post-boundary pairs (images 4-5, 5-6, and 4-6), and (3) cross-boundary pairs (images 3-4, 2-5, and 1-6). In trauma clips, this boundary corresponded to the transition from the neutral to the aversive event, whereas neutral clips followed the same temporal sampling structure without aversive content. This design allowed us to assess whether sleep differentially affected temporal memory for events occurring before, after, or across an emotional boundary.

### Functional localizer task

To aid decoding of emotional memory reactivation during sleep, we designed an emotion functional localizer task at the very end of each of the two sessions to establish the neural patterns of aversive vs. neutral processing. Participants viewed a new collection of negative and neutral images obtained from affective picture databases. The images were categorized into two subtypes based on the content: people and scenes, to be consistent with the content structure of the previously viewed traumatic and neutral film clips. Each session included 60 negative and 60 neutral images, with an equal distribution across subcategories (i.e., 30 people and 30 scene images per category). Each trial began with a jittered fixation cross (0.8-1.2 s), followed by an image presented for 3 s. To maintain attention throughout the task, participants performed a one-back working memory task. Specifically, 20% of the images were presented consecutively, and participants were instructed to press the space bar whenever they identified a repeated image.

### EEG recording

EEG was recorded using a 64-channel EEG cap with an eego amplifier (ANT Neuro, Netherland), with electrodes arranged according to the international 10-20 system and referenced online to CPz. Two electrooculogram (EOG) electrodes were placed diagonally around the eyes—one below the left eye and one above the right eye—to monitor horizontal and vertical eye movements. Two bipolar electromyogram (EMG) electrodes were positioned on the chin to record muscle activity. EEG signals were bandpass filtered online between 0.5 and 30 Hz for monitoring purposes and sampled at 500 Hz.

## Data analysis

### EEG preprocessing

Offline EEG preprocessing was performed using MNE-Python (v 1.8.0). First, the data were downsampled to 200 Hz. Next, a bandpass filter of 0.5-40 Hz and a 50 Hz notch filter were applied to remove slow drifts and line noise. The EEG signals were then re-referenced to the average of the M1 and M2 electrodes. Localizer and sleep data from both experimental sessions were segmented into epochs. For the localizer data, epochs were extracted from -1 to +3s relative to stimulus onset. These data were additionally subjected to independent component analysis (ICA), and components associated with eye blinks and eye movements were identified and removed. For the sleep data, the detection of sleep events (e.g., SOs and REMs) and the corresponding epoching procedures are described in Sleep event detection. Finally, bad channels were visually identified and interpolated, and artifact-contaminated epochs were manually rejected based on visual inspection.

### Sleep staging analysis

Sleep staging was performed using the Yet Another Spindle Algorithm (YASA, v 0.6.5), a machine learning-based sleep staging algorithm implemented in Python^80^. Raw overnight EEG data were re-referenced to FPz, following YASA’s recommended configuration. The algorithm was applied using data from the C4 electrode (or C3 if C4 was identified as a bad channel), along with EOG and EMG channels. The automatically generated hypnograms were reviewed by a trained experimenter and manually corrected as needed in accordance with standard sleep scoring guidelines.

### Sleep event detection

Slow oscillations (SOs) and sleep spindles were detected on the Cz electrode using the YASA toolbox, consistent with the central scalp topography of these oscillations^16,81^. SO and spindle analyses were restricted to N2 and N3 NREM sleep. For SO detection, the EEG signal was band-pass filtered between 0.3 and 1.5 Hz. Zero-crossings were identified, and candidate SOs were selected if they exceeded the 75th percentile of the amplitude distribution. SO-locked epochs were extracted from −2.5 to +2.5 s relative to the SO down-state negative peak. For fast spindles, the EEG was band-pass filtered between 11 and 16 Hz. The root-mean-square (RMS) amplitude was computed in sliding 300-ms windows (step 100 ms). Spindles were marked when the RMS exceeded the mean +1.5 SD of the remaining distribution. Adjacent candidates separated by less than 0.5 s were merged, and only events lasting 0.5-2 s were retained. Spindle-locked epochs were extracted from −2.5 to +2.5 s relative to the spindle peak for subsequent analyses.

To identify SO-spindle complexes, each detected SO was checked for the occurrence of a spindle within 1.5 s after its trough^16^. Matching events were classified as SO-spindle complexes. The corresponding Cz EEG signal was extracted from -2.5 s to +2.5 s around the SO trough. To further quantify SO-spindle coupling, each epoch was filtered in the SO band, 0.3-1.25 Hz, and spindle band, 11-16 Hz, using zero-phase Butterworth filters. The Hilbert transform was applied to obtain the instantaneous SO phase and spindle-band amplitude envelope. For each complex, the time point of maximal spindle-band amplitude within the detected spindle event was identified, and the corresponding SO phase angle was extracted. For each participant and session, three coupling metrics were computed: (1) the preferred coupling phase, defined as the circular mean of phase angles across all SO-spindle complexes; (2) coupling strength, quantified as the mean vector length, with higher values indicating stronger phase concentration of spindle peaks around the SO cycle; and (3) the SO-spindle coupling percentage, calculated as the number of SO-spindle complexes divided by the total number of detected spindles ^81^. The non-uniformity of preferred phases was tested using Rayleigh’s test. Between-session differences in coupling strength and coupling percentage were assessed with paired-samples t-tests, whereas between-session differences in preferred phase were tested by comparing the circular phase difference against zero using a one-sample circular test.

To identify rapid eye movements (REMs), bipolar EOG derivations were created by subtracting the left EOG from M2 and the right EOG from M1. Signals were band-pass filtered (0.5-40 Hz) and notch-filtered at 50 Hz to remove line noise. Only manually scored REM sleep epochs were analyzed. REMs were detected based on the following criteria: peak-to-peak amplitude 50-325 µV, duration 0.3-1.2 s, relative prominence ≥0.8, and dominant frequency 0.5-5 Hz^80,82,83^. Detected REMs were time-locked to their maximum peak, and phasic REMs EEG epochs (-2.5 s to +2.5 s around the peak) were extracted. To provide a control condition matched to the phasic REM events, tonic REM windows were identified as follows. First, detected REMs were temporally clustered: consecutive REMs separated by less than 5 s were grouped into the same phasic cluster. A ±5 s exclusion window was then placed around each cluster to remove phasic activity from the tonic REM candidate pool. Within the remaining REM sleep periods, continuous intervals lasting at least 30 s were identified. From these intervals, 5 s windows were selected as tonic REM candidates if they (i) were entirely contained within REM sleep, (ii) did not overlap any exclusion window, and (iii) were not adjacent to sleep-stage transitions. Finally, a random subset of tonic REM windows, matched in number to the detected phasic REM events per participant, was retained. EEG epochs from −2.5 to +2.5 s around the center of each tonic REM window were then extracted for subsequent analyses.

In addition, to characterize REM sleep at the macro level, we quantified phasic REM percentage, tonic REM percentage, and REM density. Because no unified criterion exists for defining phasic REM periods, we adopted a relatively lenient epoch-based definition to preserve eye-movement-adjacent signal. Specifically, each 30 s REM epoch was classified as a phasic period if it contained at least two detected REMs, and as a tonic period otherwise^37^. Phasic REM percentage was defined as the total duration of phasic REM periods divided by total REM sleep duration; tonic REM percentage was calculated analogously. REM density was defined as the number of detected REMs per minute of REM sleep.

### Time-frequency analyses

Time-frequency analyses were performed separately for each sleep-event type and session using Morlet wavelets in MNE-Python. Power was estimated from 1 to 30 Hz in 1 Hz steps, with the number of cycles set to frequency/2 and constrained between 1 and 5 cycles. For each channel and frequency bin, power was z-scored across all retained epochs and all time points within the full epoch window (-2.5 to +2.5 s), and then averaged across epochs to obtain participant-level time-frequency maps. The extended epoch duration was chosen to provide sufficient temporal context for low-frequency activity within the time window of interest (-1.5 to +1.5 s) while minimizing edge artifacts.

### Multivariate analysis

Multivariate classification of EEG data was performed in Python using a linear support vector machine (SVM) classifier implemented with scikit-learn, following the general decoding framework of NeuroRA^84^. For the wakeful functional localizer, EEG signals were z-scored across trials separately within each experimental session, and trials from both sessions were then collapsed within each participant. Temporal generalization decoding was conducted using a sliding-window approach with a 200 ms window and a 25 ms step size. Within each window, EEG activity was averaged across time samples to generate one feature vector per trial. To improve the signal-to-noise ratio, EEG responses from five randomly selected trials within the same valence category were averaged before classification. Classification was performed using fivefold cross-validation. Within each fold, a classifier was trained at each training-time window and tested across all testing-time windows, yielding a two-dimensional temporal generalization matrix of decoding accuracy. The diagonal of this matrix corresponds to conventional time-resolved decoding, whereas off-diagonal elements reflect temporal generalization across different time points^85^. Principal component analysis (PCA) was fitted on the training trials at each training-time window and applied to the held-out trials across testing-time windows, with the first 30 principal components retained as classification features. A linear SVM was trained to discriminate aversive from neutral trials. Classification performance was quantified as decoding accuracy, with the chance level set at 0.5. The decoding procedure was repeated 10 times, and decoding accuracy was averaged across folds and iterations for each participant.

To further examine whether emotional representations identified during wakefulness were reactivated during sleep, we used an analogous temporal generalization approach. For each participant, localizer and sleep-event EEG data were z-scored across trials independently and collapsed across sessions. Principal component analysis (PCA) was then applied to the pooled wake-sleep dataset, and the first 30 principal components were retained to define a common representational space for transfer decoding^16^. To address trial-number imbalance between wake and sleep data before PCA, sleep-event trials were randomly subsampled to match the number of localizer trials when they exceeded it; otherwise, all available sleep trials were retained. This procedure balanced the two datasets while preserving as much sleep data as possible. Temporal generalization analysis was then performed using a sliding-window approach with a 200 ms window and a 25 ms step size. Within each window, EEG activity was averaged across time samples and used as classification features. A linear SVM classifier was trained at each time window of the wakeful localizer data to discriminate aversive from neutral trials, and was then tested at each time window of the sleep-event data. This procedure yielded a temporal generalization matrix reflecting the extent to which wakeful valence-related neural patterns generalized to sleep-event activity. Because the training and testing datasets were independent, cross-validation was not required. Classification performance was quantified as accuracy, with the chance level set at 0.5. The decoding procedure was repeated 10 times, and decoding accuracy was averaged across iterations for each participant. Group-level statistical inference was performed using a nonparametric cluster-based permutation test with 1,000 permutations to correct for multiple comparisons across the temporal generalization matrix, as implemented in NeuroRA^84^. Accuracy values were compared against chance level at each train-test time point using a one-sided cluster-based permutation test. Clusters were formed from adjacent samples exceeding the cluster-forming threshold of *p* < .05, and cluster-level significance was assessed using 1,000 permutations with a corrected threshold of *p* < .05.

To assess whether phasic REM decoding was purely driven by eye-movement-related activity, we performed an ICA-based control analysis. Independent component analysis was applied to the EEG data, and components dominated by ocular activity were identified based on their spatial topographies, time courses, and correspondence with EOG activity. We then reconstructed two complementary datasets: one in which ocular-dominated components were removed, and one containing only the ocular-dominated components. The same localizer-to-sleep decoding procedure was repeated on both datasets. This analysis tested whether phasic REM decoding persisted after removal of eye-movement-related components and whether ocular-dominated activity alone was sufficient to support above-chance classification. To further examine whether phasic REMs reactivation depended on activity in a specific frequency band, we conducted a bandstop control analysis. For each canonical frequency band, both wakeful localizer data and sleep event data were filtered using the same bandstop procedure to selectively remove that frequency range while preserving the remaining broadband EEG signal. We then repeated the full decoding pipeline for each bandstop dataset to assess whether any individual frequency band was necessary for the phasic REM decoding effect.

### Statistics

Behavioral performance was analyzed using repeated-measures ANOVAs with session (aversive vs. neutral) and time (T0–T3, or pre-vs. post-sleep, depending on the analysis) as within-subject factors. Greenhouse–Geisser corrections were applied when the sphericity assumption was violated, and corrected degrees of freedom are reported accordingly. Significant interactions were followed up with simple-effects analyses using paired-samples t-tests. Partial eta-squared 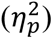 is reported as a measure of effect size and Cohen’s d are reported as measures of effect size for ANOVAs and t-tests, respectively. In control analyses, session order was additionally included as a between-subjects factor and yielded the same pattern of results. For sleep data, sleep macrostructure and microstructure were compared between sessions using paired-samples t-tests, or Wilcoxon signed-rank tests when normality assumptions were violated. To examine associations between sleep-related neural measures and behavioral outcomes, we used different statistical approaches depending on the nature of the neural metric. For conventional sleep features, including SO-spindle coupling percentage, we used linear mixed-effects models with post-sleep performance as the dependent variable: post-sleep performance ∼ session × sleep metric + pre-sleep performance + (1 | participant). Sleep metrics and pre-sleep performance were grand-mean centered prior to model fitting to facilitate interpretation of the main effects and interaction terms. For decoding-based analyses, mean decoding accuracy was extracted from statistically significant clusters for each participant. Because the temporal generalization classifier was trained to discriminate aversive from neutral stimuli and then applied to the aversive and neutral sleep events, decoding accuracy provided a participant-level index of valence-specific reactivation during sleep. We therefore related reactivation strength to differential overnight behavioral changes between the two sessions. Specifically, behavioral difference scores were first computed as the aversive minus neutral session difference at each time point, and overnight change was then calculated as the post-sleep difference minus the pre-sleep difference. Pearson correlations were used when normality assumptions were met, and Spearman rank correlations were used otherwise. Unless otherwise specified, statistical significance was set at *p* < .05.

## Supporting information

This document contains supplementary tables (1-14), figures (1-6), and notes (1-3) accompanying the main text.

## Acknowledgment

We thank Qiongyue Zhang for her assistance in data collection. The research was supported by the Ministry of Science and Technology of China STI2030-Major Projects (No. 2022ZD0214100), National Natural Science Foundation of China (No. 32171056), General Research Fund (No. 17614922) of Hong Kong Research Grants Council, and The University of Hong Kong URC Internal Research Grant to X. H..

## Author contribution

Y.Z.: conceptualization, investigation, formal analysis, software, methodology, writing – original draft, writing – review & editing, and visualization; Z.Y.: conceptualization, validation, writing – review & editing; D.C. and T.X.: conceptualization, methodology, and writing –review & editing; L.Z. : investigation and writing – review & editing; A.L.: writing – review & editing; X.H.: conceptualization, writing - original draft, writing - review & editing, supervision, project administration, and funding acquisition.

## Data and code availability

Pre-processed data and analysis scripts will be made available on the Open Science Framework (OSF) upon publication (https://osf.io/4j76c/overview?view_only=54dde3466a3949439a437307205da7e2).

## Reference

Kensinger, E. A. & Ford, J. H. Retrieval of emotional events from memory. Annu. Rev. Psychol. 71, 251–272 (2020).

LaBar, K. S. & Cabeza, R. Cognitive neuroscience of emotional memory. Nat Rev Neurosci 7, 54–64 (2006).

Iyadurai, L. et al. Intrusive memories of trauma: A target for research bridging cognitive science and its clinical application. Clinical Psychology Review 69, 67–82 (2019).

Lissek, S. & van Meurs, B. Learning models of PTSD: theoretical accounts and psychobiological evidence. International Journal of Psychophysiology 98, 594–605 (2015).

Sep, M. S. C., Geuze, E. & Joëls, M. Impaired learning, memory, and extinction in posttraumatic stress disorder: translational meta-analysis of clinical and preclinical studies. Translational Psychiatry 13, 376 (2023).

Cohen, R. T. & Kahana, M. J. A memory-based theory of emotional disorders. Psychological Review 129, 742–776 (2022).

Cabrera, Y. et al. Overnight neuronal plasticity and adaptation to emotional distress. Nat. Rev. Neurosci. 25, 253–271 (2024).

Lutz, N. D., Harkotte, M. & Born, J. Sleep’s contribution to memory formation. Physiological Reviews 106, 363–483 (2026).

Lewis, P. A. & Abdellahi, M. E. A. Could sleep engineering be used to combat PTSD and depression? PLOS Biology https://doi.org/10.1371/journal.pbio.3003633 (2026) doi:10.1371/journal.pbio.3003633.

Paller, K. A., Creery, J. D. & Schechtman, E. Memory and sleep: how sleep cognition can change the waking mind for the better. Annual Review of Psychology 72, 123–150 (2021).

Xia, T. & Hu, X. Memory editing during sleep: mechanisms, clinical applications, and technological innovations. Trends in Cognitive Sciences S1364661325002098 (2025) doi:10.1016/j.tics.2025.07.010.

Cunningham, T. J., Stickgold, R. & Kensinger, E. A. Investigating the effects of sleep and sleep loss on the different stages of episodic emotional memory: A narrative review and guide to the future. Frontiers in Behavioral Neuroscience https://doi.org/10.3389/fnbeh.2022.910317 (2022) doi:10.3389/fnbeh.2022.910317.

Yuksel, C. et al. Both slow wave and rapid eye movement sleep contribute to emotional memory consolidation. Communications Biology 8, 1–10 (2025).

Brodt, S., Inostroza, M., Niethard, N. & Born, J. Sleep—A brain-state serving systems memory consolidation. Neuron 111, 1050–1075 (2023).

Klinzing, J. G., Niethard, N. & Born, J. Mechanisms of systems memory consolidation during sleep. Nat Neurosci 22, 1598–1610 (2019).

Schreiner, T., Petzka, M., Tobias Staudigl & Staresina, B. P. Endogenous memory reactivation during sleep in humans is clocked by slow oscillation-spindle complexes. Nat Commun 12, 3112 (2021).

Schreiner, T. et al. Spindle-locked ripples mediate memory reactivation during human NREM sleep. Nat Commun 15, 5249 (2024).

Staresina, B. P. Coupled sleep rhythms for memory consolidation. Trends in Cognitive Sciences S1364661324000299 (2024) doi:10.1016/j.tics.2024.02.002.

Chen, Z. S. & Wilson, M. A. How our understanding of memory replay evolves. Journal of Neurophysiology 129, 552–580 (2023).

Cross, Z. R. et al. Slow oscillation–spindle coupling predicts sequence-based language learning. J. Neurosci. 45, e2193232024 (2025).

Diamond, N. B. et al. Sleep selectively and durably enhances memory for the sequence of real-world experiences. Nature Human Behaviour 9, 746–757 (2025).

Denis, D., Sanders, K. E. G., Kensinger, E. A. & Payne, J. D. Sleep preferentially consolidates negative aspects of human memory: Well-powered evidence from two large online experiments. Proc. Natl. Acad. Sci. U.S.A. 119, e2202657119 (2022).

Goldstein, A. N. & Walker, M. P. The role of sleep in emotional brain function. Annu. Rev. Clin. Psychol. 10, 679–708 (2014).

Lehmann, M., Schreiner, T., Seifritz, E. & Rasch, B. Emotional arousal modulates oscillatory correlates of targeted memory reactivation during NREM, but not REM sleep. Sci Rep 6, 39229 (2016).

Ben Simon, E., Rossi, A., Harvey, A. G. & Walker, M. P. Overanxious and underslept. Nat Hum Behav 4, 100–110 (2019).

Cairney, S. A., Durrant, S. J., Jackson, R. & Lewis, P. A. Sleep spindles provide indirect support to the consolidation of emotional encoding contexts. Neuropsychologia 63, 285–292 (2014).

Denis, D. et al. Slow oscillation–sleep spindle coupling is associated with expectancy measures of fear extinction retention in trauma-exposed individuals. Biological Psychiatry: Cognitive Neuroscience and Neuroimaging S2451902225002575 (2025) doi:10.1016/j.bpsc.2025.08.009.

Natraj, N. et al. Sleep spindles favor emotion regulation over memory consolidation of stressors in posttraumatic stress disorder. Biological Psychiatry: Cognitive Neuroscience and Neuroimaging S2451902223000459 (2023) doi:10.1016/j.bpsc.2023.02.007.

Rodheim, K., Kainec, K., Noh, E., Jones, B. & Spencer, R. M. C. Emotional memory consolidation during sleep is associated with slow oscillation–spindle coupling strength in young and older adults. Learn. Mem. 30, 237–244 (2023).

Greco, V. et al. Disarming emotional memories using targeted memory reactivation during rapid eye movement sleep. Imaging Neurosci (Camb) 3, IMAG.a.924 (2025).

Hutchison, I. C. et al. Targeted memory reactivation in REM but not SWS selectively reduces arousal responses. Commun Biol 4, 1–6 (2021).

Nishida, M., Pearsall, J., Buckner, R. L. & Walker, M. P. REM sleep, prefrontal theta, and the consolidation of human emotional memory. Cerebral Cortex 19, 1158–1166 (2009).

van der Helm, E. et al. REM sleep depotentiates amygdala activity to previous emotional experiences. Current Biology 21, 2029–2032 (2011).

Walker, M. P. & van der Helm, E. Overnight therapy? The role of sleep in emotional brain processing. Psychological Bulletin 135, 731–748 (2009).

Wassing, R. et al. Restless REM sleep impedes overnight amygdala adaptation. Current Biology 29, 2351–2358.e4 (2019).

Zeng, S. et al. Impaired emotional memory dissipation in insomnia disorder. Psychological Medicine 55, (2025).

Simor, P., Van Der Wijk, G., Gombos, F. & Kovács, I. The paradox of rapid eye movement sleep in the light of oscillatory activity and cortical synchronization during phasic and tonic microstates. NeuroImage 202, 116066 (2019).

Simor, P., van der Wijk, G., Nobili, L. & Peigneux, P. The microstructure of REM sleep: Why phasic and tonic? Sleep Medicine Reviews 52, 101305 (2020).

Corsi-Cabrera, M. et al. Human amygdala activation during rapid eye movements of rapid eye movement sleep: an intracranial study. Journal of Sleep Research 25, 576–582 (2016).

Maranci, J.-B. et al. Eye movement patterns correlate with overt emotional behaviours in rapid eye movement sleep. Scientific Reports 12, 1770 (2022).

Meshreky, K. M. & Lewis, P. A. Do eye movements in REM sleep play a role in overnight emotional processing? Neuropsychologia 215, 109169 (2025).

Stuart, K. & Conduit, R. Auditory inhibition of rapid eye movements and dream recall from REM sleep. Sleep 32, 399–408 (2009).

Cairney, S. A., Durrant, S. J., Power, R. & Lewis, P. A. Complementary roles of slow-wave sleep and rapid eye movement sleep in emotional memory consolidation. Cerebral Cortex 25, 1565–1575 (2015).

Tabarak, S. et al. Temporal dynamics of negative emotional memory reprocessing during sleep. Transl Psychiatry 14, 1–10 (2024).

Cunningham, T. J. & Payne, J. D. Emotional memory consolidation during sleep. in Cognitive neuroscience of memory consolidation (eds Axmacher, N. & Rasch, B.) 133–159 (Springer International Publishing, Cham, 2017). doi:10.1007/978-3-319-45066-7_9.

Tempesta, D., Socci, V., De Gennaro, L. & Ferrara, M. Sleep and emotional processing. Sleep Medicine Reviews 40, 183–195 (2018).

Holmes, E. A. & Bourne, C. Inducing and modulating intrusive emotional memories: A review of the trauma film paradigm. Acta Psychologica 127, 553–566 (2008).

James, E. L. et al. The trauma film paradigm as an experimental psychopathology model of psychological trauma: intrusive memories and beyond. Clinical Psychology Review 47, 106– 142 (2016).

Varma, M. M. et al. A systematic review and meta-analysis of experimental methods for modulating intrusive memories following lab-analogue trauma exposure in non-clinical populations. Nat Hum Behav 8, 1968–1987 (2024).

McClay, M., Sachs, M. E. & Clewett, D. Dynamic emotional states shape the episodic structure of memory. Nat Commun 14, 6533 (2023).

Petrucci, A. S. & Palombo, D. J. A matter of time: how does emotion influence temporal aspects of remembering? Cognition and Emotion 35, 1499–1515 (2021).

Wang, J., Tambini, A. & Lapate, R. C. The tie that binds: temporal coding and adaptive emotion. Trends in Cognitive Sciences 26, 1103–1118 (2022).

Schönauer, M. et al. Decoding material-specific memory reprocessing during sleep in humans. Nat Commun 8, 15404 (2017).

LaLumiere, R. T., McGaugh, J. L. & McIntyre, C. K. Emotional modulation of learning and memory: pharmacological implications. Pharmacological Reviews 69, 236–255 (2017).

McGaugh, J. L. The amygdala modulates the consolidation of memories of emotionally arousing experiences. Annual Review of Neuroscience 27, 1–28 (2004).

Kensinger, E. A. Remembering the details: effects of emotion. Emotion Review 1, 99–113 (2009).

Wang, J. & Lapate, R. C. Emotional state dynamics impacts temporal memory. Cognition and Emotion 39, 136–155 (2025).

Denis, D. et al. Sleep spindles preferentially consolidate weakly encoded memories. J Neurosci 41, 4088–4099 (2021).

Denis, D. & Payne, J. D. Targeted memory reactivation during non-rapid eye movement sleep enhances neutral, but not negative, components of memory. eNeuro ENEURO.0285-23.2024 (2024) doi:10.1523/ENEURO.0285-23.2024.

Liu, J. et al. Item-specific neural representations during human sleep support long-term memory. PLOS Biology 21, e3002399 (2023).

Schapiro, A. C., McDevitt, E. A., Rogers, T. T., Mednick, S. C. & Norman, K. A. Human hippocampal replay during rest prioritizes weakly learned information and predicts memory performance. Nat Commun 9, 3920 (2018).

Rosenblum, Y. et al. Aperiodic neural activity distinguishes between phasic and tonic REM sleep. Journal of Sleep Research 34, e14439 (2025).

Andrillon, T., Nir, Y., Cirelli, C., Tononi, G. & Fried, I. Single-neuron activity and eye movements during human REM sleep and awake vision. Nature Communications 6, 7884 (2015).

Miyauchi, S., Misaki, M., Kan, S., Fukunaga, T. & Koike, T. Human brain activity time-locked to rapid eye movements during REM sleep. Experimental Brain Research 192, 657– 667 (2008).

Chen, Z., et al. Interpreting human sleep activity through neural contrastive learning. Neuron https://www.cell.com/neuron/abstract/S0896-6273(26)00219-9 (2026).

Buysse, D. J., Reynolds, C. F., Monk, T. H., Berman, S. R. & Kupfer, D. J. The pittsburgh sleep quality index: a new instrument for psychiatric practice and research. Psychiatry Research 28, 193–213 (1989).

Beck, A. T., Steer, R. A. & Brown, G. Manual for the Beck Depression Inventory-II. (Psychological Corporation, San Antonio, TX, 1996). doi:10.1037/t00742-000.

Varma, M. M. & Hu, X. Prosocial behaviour reduces unwanted intrusions of experimental traumatic memories. Behaviour Research and Therapy 148, 103998 (2022).

Zeng, S., Lau, E. Y. Y., Li, S. X. & Hu, X. Sleep differentially impacts involuntary intrusions and voluntary recognitions of lab-analogue traumatic memories. Journal of Sleep Research 30, e13208 (2021).

Kobelt, M. et al. The memory trace of an intrusive trauma-analog episode. Current Biology 34, 1657–1669.e5 (2024).

Adan, A. & Almirall, H. Horne & östberg morningness-eveningness questionnaire: a reduced scale. Personality and Individual Differences 12, 241–253 (1991).

Spielberger, C. D., Gorsuch, Richard L., Lushene, R. E., Vagg, P. R., & Jacobs, Gerard A. Manual for the State-Trait Anxiety Inventory. (Consulting Psychologists Press, Palo Alto, CA, 1983). doi:10.30849/rip/ijp.v5i3.

Vrana, S. & Lauterbach, D. Prevalence of traumatic events and post-traumatic psychological symptoms in a nonclinical sample of college students. Journal of Traumatic Stress 7, 289– 302 (1994).

Wells, A. & Davies, M. I. The thought control questionnaire: a measure of individual differences in the control of unwanted thoughts. Behaviour Research and Therapy 32, 871– 878 (1994).

Davis, M. H. Measuring individual differences in empathy: evidence for a multidimensional approach. Journal of Personality and Social Psychology 44, 113–126 (1983).

Gross, J. J. & John, O. P. Individual differences in two emotion regulation processes: implications for affect, relationships, and well-being. Journal of Personality and Social Psychology 85, 348–362 (2003).

Watson, D., Clark, L. A. & Tellegen, A. Development and validation of brief measures of positive and negative affect: The PANAS scales. Journal of Personality and Social Psychology 54, 1063–1070 (1988).

James, E. L. et al. Computer game play reduces intrusive memories of experimental trauma via reconsolidation-update mechanisms. Psychol Sci 26, 1201–1215 (2015).

Vallat, R. & Walker, M. P. An open-source, high-performance tool for automated sleep staging. eLife 10, e70092 (2021).

Ng, T., Noh, E. & Spencer, R. M. Bayesian meta-analysis reveals the mechanistic role of slow oscillation-spindle coupling in sleep-dependent memory consolidation. eLife 13, RP101992 (2025).

Agarwal, R., Takeuchi, T., Laroche, S. & Gotman, J. Detection of rapid-eye movements in sleep studies. IEEE Transactions on Biomedical Engineering 52, 1390–1396 (2005).

Yetton, B. D. et al. Automatic detection of rapid eye movements (REMs): A machine learning approach. Journal of Neuroscience Methods 259, 72–82 (2016).

Lu, Z. & Ku, Y. NeuroRA: a python toolbox of representational analysis from multi-modal neural data. Front. Neuroinform. 14, 563669 (2020).

King, J.-R. & Dehaene, S. Characterizing the dynamics of mental representations: the temporal generalization method. Trends in Cognitive Sciences 18, 203–210 (2014).

