## Supplementary material for "Dissociable reactivation during NREM and REM sleep supports memory consolidation and emotional dissipation": This document contains supplementary tables (1-14), figures (1-6), and notes (1-3) accompanying the main text.

**This PDF file includes:**

Supporting text
Figures S1 to S6
Tables S1 to S14

**Supporting Text**

**Supplementary Note 1. Boundary-specific analyses of temporal-distance memory**

To determine whether the overall session × time interaction in temporal-distance memory was driven by specific part of the encoded events, we separately analyzed image pairs occurring before, after, or across the onset of aversive content, i.e., boundaries. For each part, temporal-distance error was submitted to a 2 (session: aversive, neutral) × 2 (time: pre-sleep, post-sleep) repeated-measures ANOVA. For pre-boundary pairs, neither the main effects of session and time nor their interaction reached significance (all $p$s $\geq$ .175), indicating no reliable session difference or overnight change in temporal-distance memory prior to the emotional boundary. For post-boundary pairs, there was a significant main effect of session, $F(1, 42) = 4.74$, $p = .035$, $\eta_p^2 = .101$, together with a significant session × time interaction, $F(1, 42) = 4.12$, $p = .049$, $\eta_p^2 = .089$. Before sleep, temporal-distance error was smaller for aversive than for neutral events, $t(42) = 3.22$, $p = .002$, whereas this difference was no longer significant after sleep, $t(42) = 0.87$, $p = .387$. Pre–post contrasts further showed that temporal-distance error decreased across sleep in the neutral session, $t(42) = 2.08$, $p = .043$, but remained stable in the aversive session, $t(42) = -0.28$, $p = .785$. For cross-boundary pairs, there was a significant main effect of time, $F(1, 42) = 7.08$, $p = .011$, $\eta_p^2 = .144$, indicating a more general overnight improvement in cross-boundary temporal estimates; the remaining effects were not significant (all $p$s $\geq$ .072).

**Supplementary Note 2. Trauma exposure disrupts sleep macrostructure but preserves event microstructure.**

To address our primary question concerning the respective roles of NREM and REM sleep in emotional memory reprocessing, we first characterized overnight sleep physiology by comparing sleep macrostructure and cardinal event microstructure following trauma-analogue and neutral film exposure. Trauma-analogue exposure significantly altered several aspects of sleep architecture (Supplementary Table 13). Relative to the neutral session, the aversive session showed reduced total sleep time (418.99 vs. 431.12 min, $p$ = .017) and lower sleep efficiency (88.18% vs. 90.28%, $p$ = .017), as well as prolonged sleep-onset latency (14.92 vs. 10.62 min, $p$ = .023) and delayed entry into N1, N2, and REM sleep (all $p$s < .030). Although absolute REM duration did not differ between sessions, $p$ = .184, the proportion of REM sleep was higher in the aversive session, $p$ = .032, with a corresponding reduction in the proportion of NREM sleep, $p$ = .032. In contrast to these macrostructural alterations, NREM- and REM-related event microstructures remained largely comparable across sessions, with the exception of rapid eye movement duration, which was slightly shorter in the neutral than in the aversive session, $t(42) = -2.56$, $p$ = .016 (Supplementary Table 14). Together, these findings indicate that trauma-analogue exposure disrupted global sleep continuity and shifted the REM/NREM balance, while leaving fine-grained sleep event characteristics largely intact.

**Supplementary Note 3. Boundary-specific associations between sleep measures and temporal-distance memory**

To determine whether the associations between SO-spindle coupling percentage and temporal-distance memory were specific to particular portions of the encoded events, we fit separate linear mixed-effects models for pre-boundary, post-boundary, and cross-boundary temporal-distance estimates. The association was most pronounced for cross-boundary estimates, with higher coupling predicting lower post-sleep error, $F(1, 61) = 4.92$, $p = .030$. A similar but weaker pattern emerged for post-boundary estimates, $F(1, 44.58) = 3.78$, $p = .058$, whereas no association was found for pre-boundary estimates, $F(1, 46.16) = 0.06$, $p = .802$. In contrast, the relationship with phasic REM percentage was localized primarily to post-boundary memory: higher phasic REM percentage predicted greater post-sleep error, $F(1, 48.27) = 5.89$, $p = .019$, whereas no significant associations were observed for pre-boundary or cross-boundary estimates (all $p$s $\geq .141$). Phasic REM density showed a comparable distribution across boundary conditions, although the individual effects did not reach statistical significance. Higher REM density was numerically associated with greater post-sleep error for post-boundary estimates, $F(1, 48.50) = 3.29$, $p = .076$, and cross-boundary estimates, $F(1, 61) = 3.18$, $p = .080$, but not for pre-boundary estimates, $F(1, 45.46) = 0.30$, $p = .587$

### Supplementary Figures

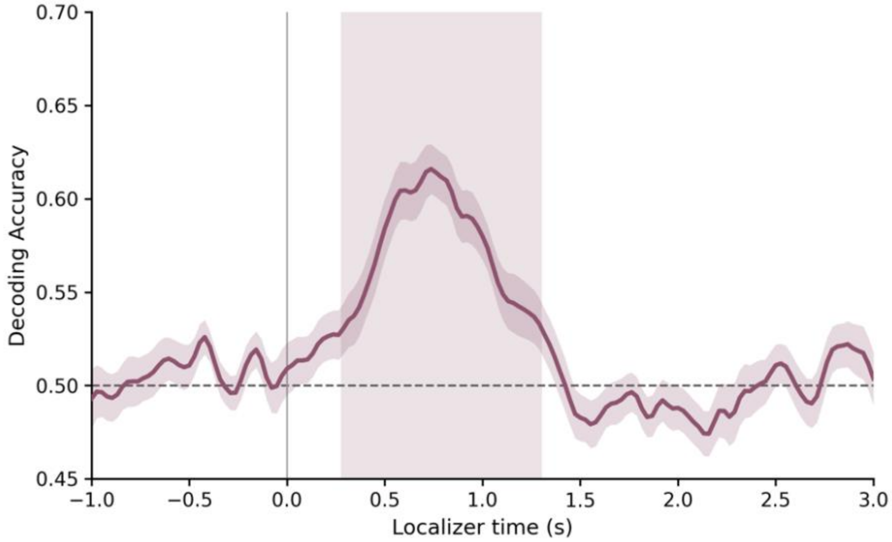


**Supplementary Fig. 1.** Time-resolved decoding successfully distinguished aversive from neutral images (diagonal of the temporal generalization matrix; training and testing at the same timepoint). Shaded areas denote significant clusters of above-chance decoding ($p_{corrected} < .05$)

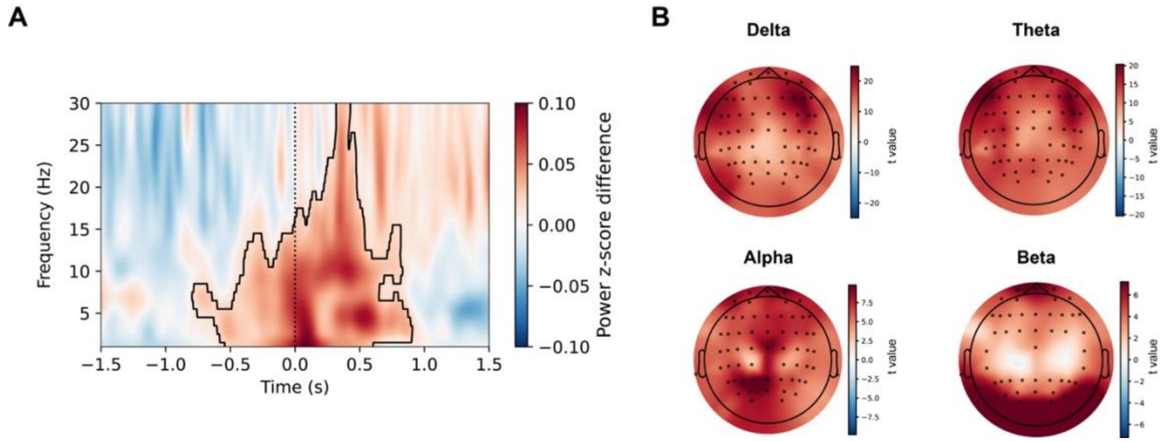


**Supplementary Fig. 2.** Spectral differences between phasic and tonic REM sleep. (A) Time–frequency contrast at Cz showing increased broadband power during phasic REM events relative to tonic REM control periods. (B) Topographic distributions of phasic-minus-tonic power differences across canonical frequency bands (−0.5 to 0.5 s relative to event onset). Phasic REM events showed widespread increases in delta, theta, alpha, and beta power relative to tonic REM control periods. Contoured areas in (a) denote significant clusters of phasic–tonic differences ($p_{corrected} < .05$); black dots in (b) mark electrodes showing significant effects.

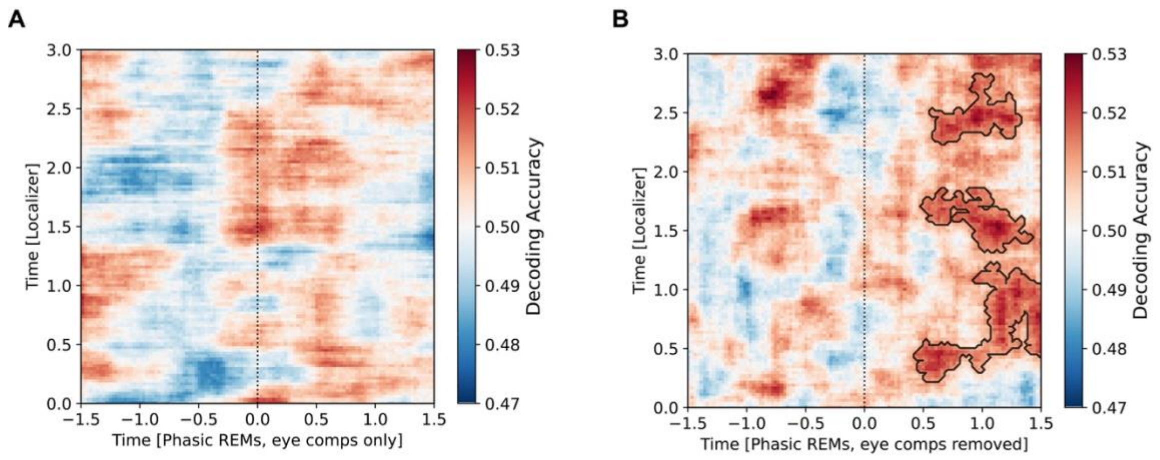


**Supplementary Fig. 3.** ICA-based control analyses for ocular components during phasic REM decoding. (A) Valence classification based only on ocular-dominated independent components did not exceed chance. (B) Significant above-chance decoding persisted after removal of ocular-dominated components, supporting a neural rather than purely ocular contribution to phasic REM valence decoding ($p_{corrected} < .05$).

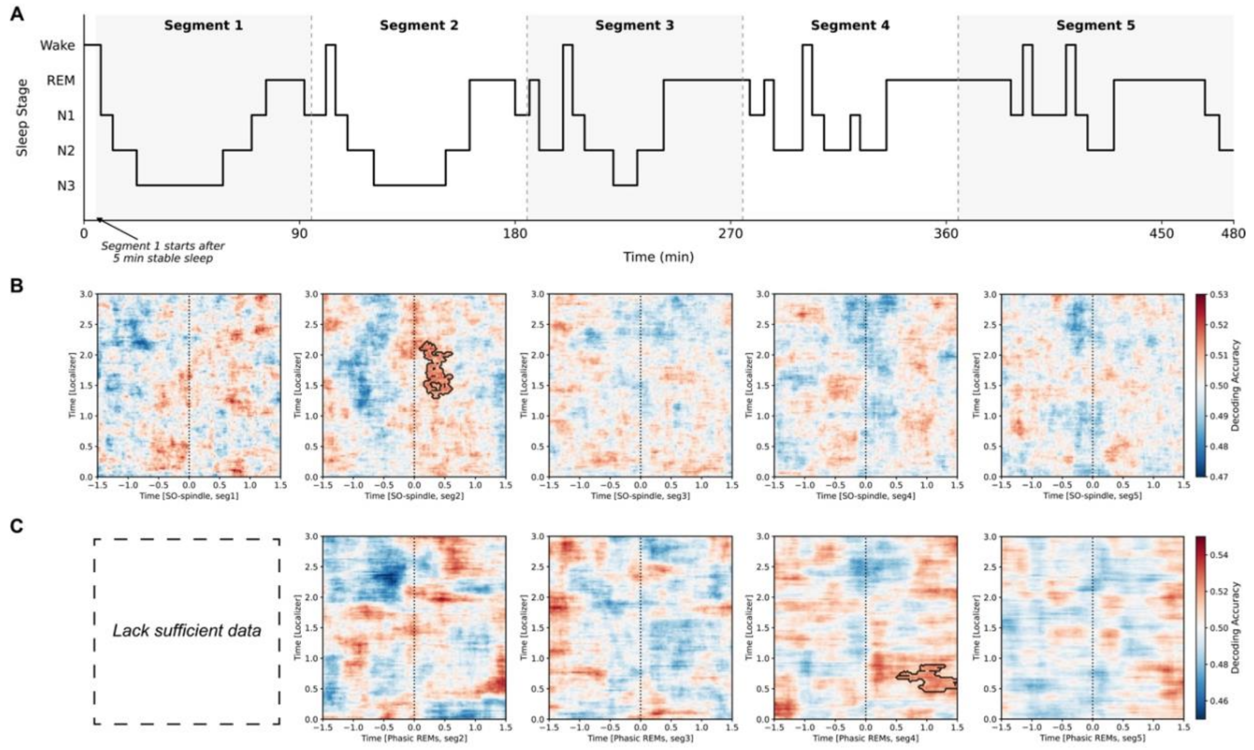


**Supplementary Fig. 4.** Segment-wise decoding of valence-specific reactivation during sleep. Sleep was divided into five temporal segments to assess whether valence-specific reactivation varied across the night. A wake-trained valence classifier was applied separately to SO-spindle complexes and phasic REM events within each segment. Significant above-chance decoding was observed for SO-spindle complexes only in segment 2 (A), whereas phasic REM showed significant above-chance decoding only in segment 4 (B), $p_{corrected} < .05$. These results suggest that emotional memory reactivation was not uniformly distributed across the night, but instead emerged in temporally specific windows for NREM SO-spindle complexes and phasic REM events.

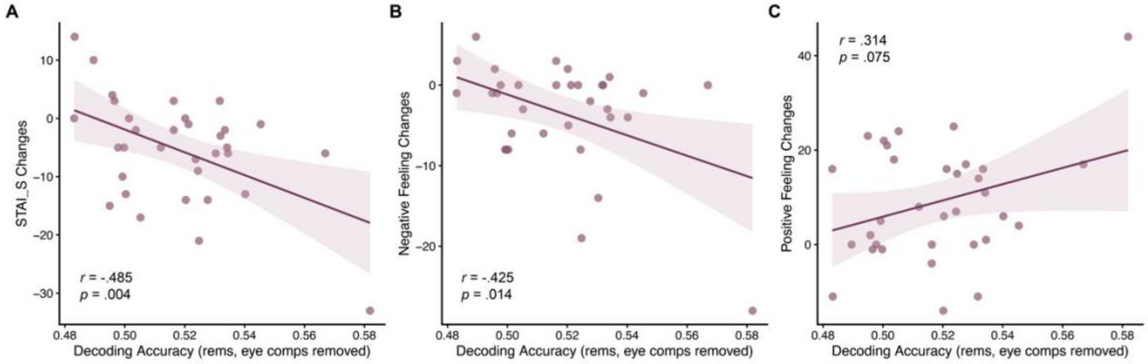


**Supplementary Fig. 5.** Reactivation-behavior associations after removal of ocular artifacts. Reactivation strength derived from ICA-corrected EEG data remained significantly associated with reductions in state anxiety and negative affect, confirming that the observed predictive effects were not driven by eye-movement artifacts.

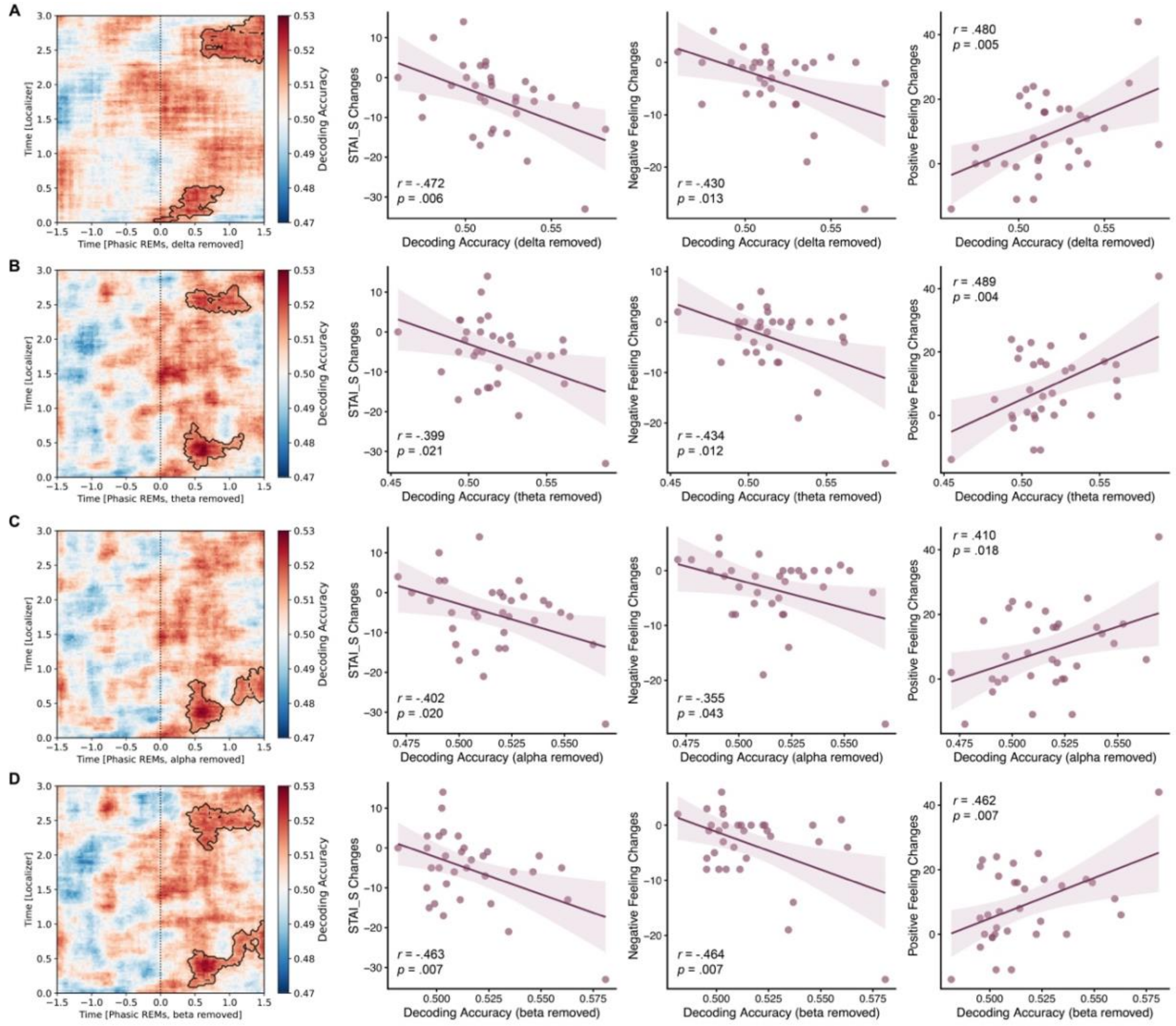


**Supplementary Fig. 6.** Phasic REM decoding is robust to single-band removal. Bandstop control analyses were performed by selectively removing individual frequency bands — (A) delta, (B) theta, (C) alpha, and (D) beta — prior to repeating the phasic REM decoding analysis. Removal of any single band did not abolish above-chance valence decoding, indicating that the decoding effect reflected broadband neural information rather than activity confined to a single frequency range ($p_{corrected} < .05$). The association between phasic REM decoding strength and overnight affective improvement was likewise preserved across all bandstop analyses.

### Supplementary Tables

| Variable | Contrast | Time | *df* | *t* | *p* |
|---|---|---|---|---|---|
| State anxiety | Aversive-Neutral | T0 | 42 | -0.18221 | 0.856 |
| State anxiety | Aversive-Neutral | T1 | 42 | 6.754265 | < .001 |
| State anxiety | Aversive-Neutral | T2 | 42 | 5.23526 | < .001 |
| State anxiety | Aversive-Neutral | T3 | 42 | 2.744509 | 0.009 |
| Negative affect | Aversive-Neutral | T0 | 42 | -2.0851 | 0.043 |
| Negative affect | Aversive-Neutral | T1 | 42 | 3.24736 | 0.002 |
| Negative affect | Aversive-Neutral | T2 | 42 | 4.467709 | < .001 |
| Negative affect | Aversive-Neutral | T3 | 42 | 1.05061 | 0.299 |
| Positive affect | Aversive-Neutral | T0 | 42 | -0.12501 | 0.901 |
| Positive affect | Aversive-Neutral | T1 | 42 | -6.70198 | < .001 |
| Positive affect | Aversive-Neutral | T2 | 42 | -6.61368 | < .001 |
| Positive affect | Aversive-Neutral | T3 | 42 | -0.54149 | 0.591 |

**Supplementary Table 1.** Post-hoc pairwise comparisons for state-level affect measures. Pairwise comparisons of session differences at each time point (T0-T3) and time differences within each session for state anxiety, negative affect, and positive affect.

| Variable | Contrast | Time | *df* | *t* | *p* |
|---|---|---|---|---|---|
| Valence | Aversive-Neutral | Pre | 42 | 13.6015 | < .001 |
| Valence | Aversive-Neutral | Post | 42 | 11.59521 | < .001 |
| Valence | Pre-Post | Aversive | 42 | 4.467709 | < .001 |
| Valence | Pre-Post | Neutral | 42 | 1.05061 | 0.299 |
| Arousal | Aversive-Neutral | Pre | 42 | -8.32559 | < .001 |
| Arousal | Aversive-Neutral | Post | 42 | -8.39916 | < .001 |
| Arousal | Pre-Post | Aversive | 42 | -0.12501 | 0.901 |
| Arousal | Pre-Post | Neutral | 42 | -6.70198 | < .001 |
| Traumatic | Aversive-Neutral | Pre | 42 | -10.6357 | < .001 |
| Traumatic | Aversive-Neutral | Post | 42 | -9.34502 | < .001 |
| Traumatic | Pre-Post | Aversive | 42 | -6.61368 | < .001 |
| Traumatic | Pre-Post | Neutral | 42 | -0.54149 | 0.591 |

**Supplementary Table 2.** Post-hoc pairwise comparisons for memory-specific emotional ratings. Pairwise comparisons of session differences before and after sleep and pre–post changes within each session for recalled valence, arousal, and traumatic impact.

| | Wake | N1 | N2 | N3 | REM | Total |
|---|---|---|---|---|---|---|
| Seg1 | 5.10 ± 7.07 | 1.81 ± 2.22 | 25.64 ± 10.75 | 48.76 ± 13.23 | 4.33 ± 5.48 | 85.64 ± 2.19 |
| Seg2 | 4.41 ± 5.40 | 3.40 ± 3.24 | 40.84 ± 12.23 | 20.73 ± 14.02 | 14.58 ± 7.88 | 83.90 ± 4.43 |
| Seg3 | 4.63 ± 6.43 | 3.07 ± 2.78 | 45.66 ± 15.07 | 12.37 ± 11.98 | 18.87 ± 11.06 | 84.60 ± 1.87 |
| Seg4 | 3.81 ± 4.74 | 4.16 ± 3.64 | 46.97 ± 12.33 | 6.01 ± 9.19 | 23.35 ± 9.86 | 84.30 ± 1.95 |
| Seg5 | 6.09 ± 7.73 | 5.76 ± 4.25 | 40.01 ± 18.69 | 3.93 ± 8.00 | 32.05 ± 16.00 | 87.83 ± 25.28 |

**Supplementary Table 3.** Duration of sleep stages by segment (min, mean ± SD). Note: Segments were defined as consecutive 90-minute intervals, with Segment 1 beginning at stable sleep onset (defined as at least 5 consecutive minutes of sleep)

| Event | Segment | Valid | Neutral | | Aversive | |
|---|---|---|---|---|---|---|
| | | | M | SD | M | SD |
| Solitary SOs | Seg1 | 33 | 1657.24 | 279.95 | 1591.33 | 422.01 |
| | Seg2 | 33 | 1144.91 | 258.22 | 1182.55 | 265.38 |
| | Seg3 | 33 | 956.21 | 279.92 | 1004.91 | 272.91 |
| | Seg4 | 33 | 856.15 | 280.87 | 825.09 | 298.65 |
| | Seg5 | 33 | 841.85 | 338.5 | 817.09 | 362.6 |
| Spindle | Seg1 | 33 | 146.82 | 69.67 | 148.64 | 83.33 |
| | Seg2 | 32 | 203.50 | 77.87 | 198.38 | 82.36 |
| | Seg3 | 33 | 220.91 | 89.19 | 202.09 | 84.73 |
| | Seg4 | 33 | 226.03 | 90.46 | 222.39 | 89.10 |
| | Seg5 | 32 | 206.91 | 104.01 | 192.22 | 88.13 |
| SO-Spindle complexes | Seg1 | 20 | 80.95 | 31.76 | 83.20 | 42.53 |
| | Seg2 | 27 | 101.48 | 41.01 | 91.19 | 36.65 |
| | Seg3 | 29 | 102.24 | 41.46 | 95.86 | 39.86 |
| | Seg4 | 30 | 113.67 | 51.53 | 104.57 | 47.08 |
| | Seg5 | 27 | 113.30 | 53.28 | 113.89 | 46.57 |
| Phasic REMs | Seg1 | - | - | - | - | - |
| | Seg2 | 15 | 87.33 | 41.84 | 119 | 99.51 |
| | Seg3 | 17 | 128.47 | 96.41 | 83.24 | 49.45 |
| | Seg4 | 24 | 148.42 | 94.6 | 191.38 | 140.01 |
| | Seg5 | 28 | 208.64 | 113.24 | 222.71 | 158.75 |
| Tonic REMs | Seg1 | - | - | - | - | - |
| | Seg2 | 15 | 69.60 | 31.47 | 86.13 | 56.27 |
| | Seg3 | 16 | 97.62 | 43.07 | 80.31 | 46.00 |
| | Seg4 | 24 | 115.08 | 60.03 | 141.58 | 81.37 |
| | Seg5 | 28 | 186.54 | 97.92 | 182.82 | 117.24 |

**Supplementary Table 4.** Trial counts for sleep event decoding analyses across sleep segment. Note: For robust decoding, each session required at least 30 trials (events) and data from at least 12 participants. Valid = number of participants with sufficient data.

| Sleep Event | *r* | *p* | *n* | Partial *r* | Partial *p* | Partial *df* |
|---|---|---|---|---|---|---|
| SO-spindle complexes | .010 | .955 | 33 | -.012 | .949 | 27 |
| Phasic REMs | -.084 | .643 | 33 | -.189 | .325 | 27 |
| Phasic REMs (ICA removed) | -.088 | .625 | 33 | -.143 | .459 | 27 |

**Supplementary Table 5.** Association between mean decoding accuracy within the significant cluster and overnight changes in absolute temporal distance error. Partial correlations (*r*) were calculated controlling for sleep architecture parameters (TST, SOL, REM%, and SE).

| Sleep Event | *Behavioral measures* | *r* | *p* | n | Partial *r* | Partial *p* | Partial *df* |
|---|---|---|---|---|---|---|---|
| SO-spindle complexes | Valence | .141 | .433 | 33 | .162 | .402 | 27 |
| | Arousal | .257 | .149 | 33 | .276 | .148 | 27 |
| | Traumatic | -.062 | .734 | 33 | -.101 | .603 | 27 |
| Phasic REMs | Valence | .282 | .112 | 33 | .216 | .260 | 27 |
| | Arousal | -.008 | .966 | 33 | -.075 | .699 | 27 |
| | Traumatic | .009 | .960 | 33 | .002 | .992 | 27 |
| Phasic REMs (ICA removed) | Valence | .201 | .262 | 33 | .161 | .405 | 27 |
| | Arousal | -.112 | .536 | 33 | -.190 | .322 | 27 |
| | Traumatic | .086 | .633 | 33 | .024 | .901 | 27 |

**Supplementary Table 6.** Association between mean decoding accuracy within the significant cluster and overnight changes in memory-specific emotional responses (valence, arousal and traumatic feelings). Partial correlations (*r*) were calculated controlling for sleep architecture parameters (TST, SOL, REM%, and SE).

| Sleep Event | *Behavioral measures* | *r* | *p* | n | Partial *r* | Partial *p* | Partial *df* |
|---|---|---|---|---|---|---|---|
| SO-spindle complexes | STAI_S | -.173 | .335 | 33 | -.178 | .355 | 27 |
| | Negative | -.089 | .624 | 33 | -.080 | .681 | 27 |
| | Positive | .063 | .728 | 33 | .034 | .859 | 27 |
| Phasic REMs | STAI_S | -.461 | .007 | 33 | -.566 | .001 | 27 |
| | Negative | -.468 | .006 | 33 | -.545 | .002 | 27 |
| | Positive | .436 | .011 | 33 | .526 | .003 | 27 |
| Phasic REMs (ICA removed) | STAI_S | -.485 | .004 | 33 | -.520 | .004 | 27 |
| | Negative | -.425 | .014 | 33 | -.465 | .011 | 27 |
| | Positive | .314 | .075 | 33 | .362 | .054 | 27 |

**Supplementary Table 7.** Association between mean decoding accuracy within the significant cluster and overnight changes in general emotional state (STAI_S, negative and positive affect). Partial correlations (*r*) were calculated controlling for sleep architecture parameters (TST, SOL, REM%, and SE).

| Variables | $r$ | $p$ | $p_{FDR}$ |
|---|---|---|---|
| BDI-II | .03 | .870 | 0.909 |
| STAI_T | 0.258 | 0.148 | 0.338 |
| TCQ | 0.245 | 0.170 | 0.340 |
| Distraction | 0.021 | 0.909 | 0.909 |
| Social control | 0.103 | 0.568 | 0.772 |
| Worry | 0.462 | 0.007 | 0.112 |
| Punishment | 0.149 | 0.407 | 0.651 |
| Reappraisal | -0.291 | 0.101 | 0.301 |
| IRI | 0.360 | 0.040 | 0.213 |
| Perspective taking | 0.010 | 0.579 | 0.772 |
| Fantasy | 0.053 | 0.768 | 0.909 |
| Empathic concern | 0.150 | 0.406 | 0.651 |
| Personal distress | 0.377 | 0.031 | 0.213 |
| ERQ | -0.281 | 0.113 | 0.301 |
| Reappraisal | -0.047 | 0.796 | 0.909 |
| Suppression | -0.305 | 0.084 | 0.301 |

**Supplementary Table 8.** Assessing the association between trait-like characteristics and emotional memory reactivation during phasic REMs.

| Variables | Mean ± SD (n, %) | Median | Range |
|---|---|---|---|
| Age | 24.37 ± 3.01 | 24 | [19, 31] |
| Gender | Female: 34, 79.07% | / | / |
| PSQI | 3.46 ± 1.44 | 3 | [1, 7] |
| Sleep time | 23:38:00<br>± 34.00 min | 23:30:00 | [22:00:00, 00:40:00] |
| Awake time | 08:23:00<br>± 47.97 min | 08:30:00 | [06:00:00, 10:15:00] |
| Duration | 8.03 ± 0.53 h | 8.00 | [7.00, 9.00] |
| Latency | 16.35 ± 8.29 min | 15 | [5, 30] |
| rMEQ | 2: 5, 11.63%<br>3: 34, 79.07%<br>4: 4, 9.30% | / | / |
| BDI-II | 2.86 ± 3.32 | 2 | [0, 14] |
| STAI_S | 29.54 ± 7.42 | 29 | [20, 51] |
| STAI_T | 32.95 ± 8.51 | 32 | [20, 55] |
| TEQ | 0.47 ± 1.03 | 0 | [0, 5] |
| TCQ | 65.91 ± 8.98 | 65 | [50, 86] |
| Distraction | 16.95 ± 3.15 | 17 | [10, 24] |
| Social control | 14.54 ± 2.81 | 14 | [11, 19] |
| Worry | 9.28 ± 3.37 | 9 | [6, 19] |
| Punishment | 9.61 ± 2.74 | 9 | [6, 17] |
| Reappraisal | 15.54 ± 2.76 | 16 | [10, 21] |
| IRI | 58.14 ± 10.48 | 56 | [35, 78] |
| Perspective taking | 13.88 ± 2.59 | 14 | [7, 19] |
| Fantasy | 17.95 ± 5.40 | 18 | [2, 26] |
| Empathic concern | 19.58 ± 4.77 | 20 | [5, 26] |
| Personal distress | 6.72 ± 4.58 | 6 | [0, 20] |
| ERQ | 42.44 ± 7.88 | 42 | [20, 57] |
| Reappraisal | 30.33 ± 4.86 | 31 | [15, 40] |
| Suppression | 12.12 ± 5.93 | 11 | [4, 27] |

**Supplementary Table 9.** Demographic and psychological characteristics of the participants (N = 43).

| Category | Feature | Neutral | Aversive | $t$ | $p$ |
| --- | --- | --- | --- | --- | --- |
| Content | / | 3.68 ± 0.36 | 3.68 ± 0.36 | 0.00 | 1.000 |
| Emotional feelings | Valence | 5.01 ± 0.47 | 2.76 ± 0.53 | 11.00 | < 0.001 |
| | Arousal | 2.36 ± 0.79 | 6.00 ± 0.74 | -11.65 | < 0.001 |
| | Traumatic | 1.07 ± 0.15 | 6.39 ± 0.74 | -24.42 | < 0.001 |
| Discrete emotional feelings | Sad | 1.10 ± 0.23 | 4.44 ± 0.51 | -20.59 | < 0.001 |
| | Hopeless | 1.00 ± 0.00 | 5.57 ± 1.10 | -14.45 | < 0.001 |
| | Fearful | 1.36 ± 0.39 | 5.79 ± 0.84 | -16.57 | < 0.001 |
| | Anxious | 5.61 ± 0.73 | 5.07 ± 0.88 | 1.65 | 0.114 |
| | Depressed | 4.22 ± 0.47 | 4.17 ± 0.55 | 0.27 | 0.793 |
| | Disgust | 2.14 ± 0.61 | 4.47 ± 1.81 | -4.24 | < 0.001 |
| Story | Storyline | 1.89 ± 0.36 | 2.14 ± 0.37 | -1.67 | 0.109 |
| | Event_units | 6.89 ± 0.83 | 9.79 ± 1.41 | -6.13 | < 0.001 |
| Visual | Visual_all | 2.44 ± 0.40 | 2.86 ± 0.40 | -2.53 | 0.019 |
| | Central_num | 2.70 ± 0.91 | 2.42 ± 1.42 | 0.58 | 0.569 |
| | Background_num | 2.65 ± 1.05 | 3.00 ± 1.14 | -0.78 | 0.446 |
| | Colour | 2.43 ± 0.51 | 2.50 ± 0.53 | -0.33 | 0.746 |
| | Movement | 2.26 ± 0.53 | 3.11 ± 0.54 | -3.89 | < 0.001 |
| | Frame_transitions | 2.17 ± 0.46 | 3.26 ± 0.69 | -4.58 | < 0.001 |
| Sound | Sound_all | 2.27 ± 0.35 | 2.22 ± 0.25 | 0.37 | 0.713 |
| | Speech | 2.69 ± 0.73 | 1.76 ± 0.53 | 3.55 | 0.002 |
| | Music | 1.86 ± 0.70 | 1.65 ± 0.42 | 0.88 | 0.387 |
| | Noise | 2.25 ± 0.47 | 3.25 ± 0.60 | -4.53 | < 0.001 |

**Supplementary Table 10.** Comparison of aversive and neutral film clips across semantic, emotional, and perceptual dimensions.

| Picture ID | Neutral | Aversive | *t* | *p* |
| --- | --- | --- | --- | --- |
| Pic#1 | 13.92 ± 3.09 | 13.33 ± 3.55 | 0.43 | 0.672 |
| Pic#2 | 33.58 ± 6.10 | 31.08 ± 5.90 | 1.02 | 0.318 |
| Pic#3 | 53.83 ± 7.58 | 49.42 ± 9.52 | 1.26 | 0.222 |
| Pic#4 | 73.00 ± 9.71 | 73.42 ± 10.23 | -0.10 | 0.919 |
| Pic#5 | 92.92 ± 6.96 | 92.08 ± 7.46 | 0.28 | 0.780 |
| Pic#6 | 112.42 ± 4.29 | 110.92 ± 6.30 | 0.68 | 0.504 |

**Supplementary Table 11.** Temporal sampling points (in seconds) of still frames extracted from film clips.

| Condition | Valence | | | | Arousal | | | |
|---|---|---|---|---|---|---|---|---|
| | Aversive | Neutral | *t* | *p* | Aversive | Neutral | *t* | *p* |
| People (S1) | 2.36 ± 0.60 | 4.93 ± 1.85 | -7.02 | < .001 | 6.56 ± 0.69 | 3.16 ± 1.59 | 10.41 | < .001 |
| Scene (S1) | 1.67 ± 2.09 | 5.11 ± 1.97 | -6.38 | < .001 | 4.64 ± 2.23 | 3.36 ± 1.75 | 2.42 | 0.019 |
| People (S2) | 2.16 ± 1.48 | 4.87 ± 1.92 | -5.92 | < .001 | 6.06 ± 1.62 | 3.14 ± 1.62 | 6.80 | < .001 |
| Scene (S2) | 1.53 ± 2.18 | 4.53 ± 2.31 | -5.09 | < .001 | 4.47 ± 2.24 | 3.06 ± 2.05 | 2.50 | 0.015 |

**Supplementary Table 12.** Normative ratings of valence and arousal for functional localizer stimuli across counterbalanced subsets.

| Variables | Neutral | | Aversive | | Statistics | | |
|---|---|---|---|---|---|---|---|
| | M | SD | M | SD | *t/W* | *p* | Effect size |
| TIB | 477.61 | 8.5 | 475.21 | 14.12 | 329 | 0.825 | 0.044 |
| SPT | 466.42 | 18.19 | 459.6 | 24.18 | 451 | 0.065 | 0.354 |
| WASO | 35.29 | 18.25 | 40.61 | 32.86 | 271.5 | 0.338 | -0.185 |
| TST | 431.12 | 26.89 | 418.99 | 36.35 | 461 | 0.017 | 0.463 |
| N1 | 28.47 | 17.15 | 24.15 | 10.12 | 415.5 | 0.101 | 0.319 |
| N2 | 213.72 | 30.59 | 201.96 | 29.52 | 1.86 | 0.072 | 0.309 |
| N3 | 94.39 | 24.43 | 93.92 | 19.99 | 0.14 | 0.889 | 0.023 |
| REM | 94.54 | 21.94 | 98.96 | 22.59 | -1.35 | 0.184 | -0.226 |
| NREM | 336.58 | 26.28 | 320.03 | 33.67 | 2.97 | 0.005 | 0.495 |
| SOL | 10.62 | 16.15 | 14.92 | 17.85 | 176 | 0.023 | -0.441 |
| Lat_N1 | 10.69 | 16.26 | 21.46 | 30.29 | 132.5 | 0.003 | -0.579 |
| Lat_N2 | 14.9 | 16.05 | 26.19 | 30.44 | 174 | 0.013 | -0.477 |
| Lat_N3 | 23.88 | 16.56 | 31.36 | 24.35 | 218 | 0.072 | -0.345 |
| Lat_REM | 94.53 | 29.92 | 109.14 | 36.9 | -2.4 | 0.022 | -0.4 |
| %N1 | 6.64 | 3.99 | 5.79 | 2.5 | 407 | 0.248 | 0.222 |
| %N2 | 49.61 | 6.7 | 48.13 | 5.49 | 1.21 | 0.233 | 0.202 |
| %N3 | 21.88 | 5.49 | 22.48 | 4.63 | -0.72 | 0.474 | -0.121 |
| %REM | 21.88 | 4.6 | 23.6 | 4.76 | 196 | 0.032 | -0.411 |
| %NREM | 78.12 | 4.6 | 76.4 | 4.76 | 470 | 0.032 | 0.411 |
| SE | 90.28 | 5.59 | 88.18 | 7.26 | 485 | 0.017 | 0.456 |
| SME | 92.41 | 3.97 | 91.22 | 6.8 | 404 | 0.268 | 0.213 |

**Supplementary Table 13.** Comparison of sleep macrostructure between neutral and aversive sessions. Note. Sleep parameters are presented in minutes unless otherwise specified. Time in Bed (TIB): total duration of the hypnogram; Sleep Period Time (SPT): duration from first to last period of sleep; Wake After Sleep Onset (WASO): duration of wake periods within SPT; Total Sleep Time (TST): total duration of N1 + N2 + N3 + REM sleep in SPT; N1, N2, N3 and REM: sleep stages duration. NREM = N1 + N2 + N3; Sleep Onset Latency (SOL): Latency to first epoch of any sleep; Latencies: latencies of sleep stages (N1, N2, N3, REM) from the beginning of the record; % (W, … REM): sleep stages duration expressed in percentages of TST; Sleep Efficiency (SE): TST / TIB * 100 (%); Sleep Maintenance Efficiency (SME): TST / SPT * 100 (%).

| Variables | Neutral | | Aversive | | Statistics | | |
|---|---|---|---|---|---|---|---|
| | M | SD | M | SD | *t/W* | *p* | Effect size |
| **NREM** | | | | | | | |
| Spindle count | 1004.94 | 330.64 | 946.39 | 310.18 | 1.99 | 0.055 | 0.347 |
| Spindle density | 3.28 | 1.06 | 3.23 | 1.05 | 0.78 | 0.442 | 0.136 |
| Spindle duration | 0.86 | 0.06 | 0.85 | 0.06 | 1.17 | 0.250 | 0.204 |
| Spindle amplitude | 68.37 | 8.20 | 71.01 | 17.31 | 290 | 0.872 | 0.034 |
| Spindle frequency | 12.95 | 0.37 | 12.91 | 0.38 | 0.87 | 0.390 | 0.152 |
| SO count | 6559.85 | 985.37 | 6541.55 | 1068.54 | 286 | 0.929 | 0.020 |
| SO density | 21.34 | 2.86 | 22.03 | 2.59 | 201 | 0.158 | -0.283 |
| SO duration | 1.15 | 0.03 | 1.14 | 0.02 | 334 | 0.194 | 0.265 |
| SO PTP | 53.21 | 15.64 | 52.61 | 14.40 | 315 | 0.544 | 0.123 |
| SO frequency | 0.93 | 0.02 | 0.94 | 0.02 | -1.83 | 0.077 | -0.318 |
| SO-spindle complex count | 439.70 | 197.81 | 419.94 | 195.31 | 1.03 | 0.309 | 0.180 |
| SO-spindle complex density | 1.44 | 0.65 | 1.43 | 0.65 | 0.18 | 0.861 | 0.031 |
| SO-spindle coupling phase | -0.28 | 0.26 | -0.29 | 0.27 | 0.05 | 0.958 | 0.009 |
| SO-spindle coupling strength | 0.32 | 0.07 | 0.32 | 0.07 | 262 | 0.748 | -0.066 |
| **REM** | | | | | | | |
| REM segments | 6.33 | 1.73 | 6.15 | 1.91 | 0.49 | 0.627 | 0.085 |
| Phasic REM proportion | 36.66 | 12.60 | 36.37 | 12.93 | 0.17 | 0.869 | 0.029 |
| Tonic REM proportion | 53.26 | 12.11 | 54.36 | 12.94 | -0.65 | 0.519 | -0.113 |
| Rapid eye movement count | 457.36 | 209.15 | 506.15 | 303.28 | 206 | 0.186 | -0.266 |
| Rapid eye movement density | 5.05 | 2.68 | 5.07 | 2.81 | 280 | 1.000 | -0.002 |
| Rapid eye movement duration | 0.76 | 0.03 | 0.77 | 0.03 | -2.56 | 0.016 | -0.445 |

**Supplementary Table 14.** Comparison of sleep microstructural features between the neutral and aversive sessions. SO = slow oscillation; PTP = peak-to-peak amplitude; REM = rapid eye movement. NREM microstructural features, including SOs, spindles, and SO-spindle complexes, were detected at the Cz electrode; rapid eye movements were detected from bipolar EOG derivations.
